# Human ADA2 deficiency is characterized by the absence of an intracellular hypoglycosylated form of adenosine deaminase 2

**DOI:** 10.1101/2023.10.25.564037

**Authors:** Lisa Ehlers, Marjon Wouters, Anneleen Hombrouck, Bethany Pillay, Selket Delafontaine, Giorgia Bucciol, Marco Baggio, Mariia Dzhus, Emma Francés Rabanal, Alexandra Damerau, Julia Neugebauer, Frédéric Ebstein, Maarten Jacquemyn, Lien De Somer, Rik Schrijvers, Steven Vanderschueren, David Cassiman, Marieluise Kirchner, Philipp Mertins, Mir-Farzin Mashreghi, Tilmann Kallinich, Dirk Daelemans, Patrizia Agostinis, Leen Moens, Isabelle Meyts

**Affiliations:** Department of Microbiology, Immunology and Transplantation, Laboratory for Inborn Errors of Immunity, KU Leuven, Leuven, Belgium; Department of Pediatric Respiratory Medicine, Immunology and Critical Care Medicine, Charité – Universitätsmedizin Berlin, corporate member of Freie Universität Berlin and Humboldt-Universität zu Berlin, Berlin, Germany; Berlin Institute of Health at Charité – Universitätsmedizin Berlin, Berlin, Germany; German Center for Child and Adolescent Health (DZKJ), partner site Berlin, Berlin, Germany; Deutsches Rheuma-Forschungszentrum, an Institute of the Leibniz Association, Berlin, Germany; Department of Pediatrics, University Hospitals Leuven, KU Leuven, Leuven, Belgium; Laboratory of Computational and Developmental Biology, Berlin Institute for Medical Systems Biology (BIMSB), Max-Delbrück-Centrum for Molecular Medicine in the Helmholtz Association (MDC), Berlin, Germany; Department of Rheumatology and Clinical Immunology, Charité – Universitätsmedizin Berlin, corporate member of Freie Universität Berlin and Humboldt-Universität zu Berlin, Berlin, Germany; Department of Pediatric Gastroenterology, Nephrology and Metabolic Diseases, Center of Chronically Sick Children, Charité – Universitätsmedizin Berlin, corporate member of Freie Universität Berlin and Humboldt-Universität zu Berlin, Berlin, Germany; Nantes University, CHU Nantes, CNRS, INSERM, Nantes, France; KU Leuven, Department of Microbiology, Immunology and Transplantation, Rega Institute for Medical Research, Molecular Genetics and Therapeutics in Virology and Oncology Research Group, Leuven, Belgium; Department of Pediatric Rheumatology, University Hospitals Leuven, Leuven, Belgium; Laboratory of Immunobiology, Department of Microbiology and Immunology, KU Leuven, Leuven, Belgium; Department of Microbiology, Immunology and Transplantation, Allergy and Clinical immunology Research group, KU Leuven, Leuven, Belgium; Department of General Internal Medicine, Research Department Microbiology, Immunology, and Transplantation, Laboratory of Clinical Infectious and Inflammatory Disorders, University Hospitals Leuven, Leuven, Belgium; Department of Gastroenterology-Hepatology and Metabolic Center, University Hospital Leuven, Leuven, Belgium; Core Unit Proteomics, Berlin Institute of Health at Charité - Universitätsmedizin Berlin and Max-Delbrück-Center for Molecular Medicine, Berlin, Germany; Cell Death Research & Therapy (CDRT) Lab, Department of Cellular & Molecular Medicine, Center for Cancer Biology, VIB-KU Leuven, Leuven, Belgium

**Author notes:** **Corresponding author:** Isabelle Meyts, Laboratory for Inborn Errors of Immunity and Department of Pediatrics UZ Leuven, Herestraat 49, 3000 Leuven, Belgium.

## Abstract

Human deficiency of adenosine deaminase 2 (DADA2) is an autoinflammatory disease caused by pathogenic variants in *ADA2* that lead to impaired deaminase activity. Recently, a lysosomal function of ADA2 has been proposed but an intracellular form of the protein has not yet been characterized. Here, we analyze protein expression of mutant ADA2 in human monocyte-derived macrophages from 10 DADA2 patients. We identify an intracellular low-molecular-weight (LMW) form of ADA2 that undergoes glycan trimming by α-mannosidases and is absent in DADA2 macrophages. Subcellular fractionation and immunofluorescence microscopy demonstrate that LMW-ADA2 is localized in the lysosomes. By overexpression of 34 *ADA2* variants in HEK293T and U-937 cells, we show that absence of LMW-ADA2 strongly correlates with reduced deaminase activity and predicts variant pathogenicity. In conclusion, we describe a previously unreported intracellular hypoglycosylated form of ADA2 and establish the absence of this LMW-ADA2 as a cellular characteristic of DADA2. Thereby, we introduce a protein correlate of the recently described lysosomal form of ADA2.

**Summary:** Ehlers et al. demonstrate that mutant ADA2 fails to undergo glycan processing beyond the ER leading to the absence of intracellular low-molecular-weight (LMW) ADA2 in DADA2 patients’ macrophages. LMW-ADA2 localizes to the lysosomes and can be generated from extracellular wild-type ADA2.

## Introduction

Human deficiency of adenosine deaminase 2 (DADA2) is an inborn error of immunity caused by biallelic deleterious mutations in the *ADA2* gene (*1, 2*). The disease manifests with a diverse phenotype displaying features of both autoinflammation and immunodeficiency (*3*). More than 600 missense or predicted loss-of-function variants in *ADA2* are listed in the Genome Aggregation Database (gnomAD v4.1.0) (*4*). In the Infevers registry, 116 *ADA2* variants are classified as (likely) pathogenic (*5*). Complete loss-of-function variants are more likely to cause bone marrow failure. Missense variants which account for 80% of pathogenic variants are found in patients with a primarily vasculitis phenotype as well as in those presenting with cytopenias and hypogammaglobulinemia (*6, 7*). Our group recently demonstrated that specific monoallelic *ADA2* variants can cause ADA2 deficiency through a dominant negative mechanism (*8*).

How mutations in *ADA2* give rise to this diverse phenotype, even in patients with identical genotypes, is not understood. Inflammatory symptoms of the disease typically respond well to tumor necrosis factor-alpha inhibitors (TNFi) (*9*). Insufficient resolution of bone marrow failure in response to this treatment however often necessitates hematopoietic stem cell transplantation and disease lethality still is 8% (*10*). To develop improved treatment options, we require a better understanding of the pathomechanisms underlying the immunological and clinical phenotype of DADA2.

ADA2 is a 59-kD glycoprotein with a signal peptide that mediates secretion of the protein via the endoplasmic reticulum (ER), and exhibits extracellular adenosine deaminase activity as a homodimer (*11*). The protein has four N-glycosylation sites and insufficient glycosylation leads to ER retention and intracellular aggregate formation of ADA2 transfected into HEK293T cells (*12, 13*). Impaired protein secretion and absent serum ADA2 enzyme activity have been established as characteristics shared by the majority of pathogenic *ADA2* variants (*14*). Mechanistic approaches to the pathophysiology of ADA2-deficient cells have therefore been based on impaired extracellular adenosine deamination driving inflammation (*15, 16*). This assumption fails to acknowledge the weak affinity of ADA2 for adenosine and the presence of functional ADA1 in DADA2 patients (*17*). Research into alternative functions of the ADA2 protein and DADA2-causing variants is therefore warranted. Biochemical characterization of pathogenic *ADA2* variants has until now mostly been performed in a HEK293T overexpression system with supraphysiological intracellular levels of ADA2 while expression of the mutant protein has been shown to be minimal in DADA2 CD14^+^ monocytes (*14, 18, 19*). Disease-associated *ADA2* variants have to date hardly been studied in their physiological milieu.

In this study, we analyze trafficking and glycosylation of endogenous ADA2 expressed in primary human macrophages from healthy donors and DADA2 patients. We provide evidence of an intracellular glycoform of ADA2 that results from glycan trimming by Golgi α-mannosidases and show that DADA2 macrophages do not express this low-molecular-weight-form of ADA2. Absence of LMW-ADA2 correlates with reduced ADA2 enzyme activity and serves as a biomarker of variant pathogenicity.

## Results

### *In vitro* macrophage differentiation partially restores ADA2 protein expression in DADA2 monocytes

Of the circulating leukocytes, monocytes express ADA2 most abundantly. We analyzed ADA2 protein expression and *ADA2* mRNA levels in peripheral blood monocytes of 10 DADA2 patients and three heterozygous carriers (**Table 1** and **Fig. S1A+B**). CD14^+^ monocytes from DADA2 patients showed very low or absent ADA2 protein expression despite normal *ADA2* mRNA levels in patients with missense mutations which account for 80% of pathogenic variants found in DADA2 patients (**Fig. 1A** and **Fig. S2A**). Upon macrophage differentiation, we observed an increase in mutant ADA2 protein expression in the patient cells while ADA2 protein expression was reduced in healthy control (HC) human monocyte-derived macrophages (HMDM) (**Fig. 1B**). Successful differentiation of DADA2 HMDM was achieved both with granulocyte–macrophage colony-stimulating factor (GM-CSF) and macrophage colony-stimulating factor (M-CSF) (**Fig. S2B**), which confirms the reported restoration of macrophage differentiation of monocytes from DADA2 patients under treatment with TNFi (*9*). Interestingly, western blotting of pathogenic ADA2 protein variants expressed in DADA2 HMDM revealed the presence of only a high-molecular-weight (HMW) band of around 60 kD compared to HC HMDM displaying two bands at 60 and 57 kD, with the low-molecular-weight (LMW) band being more pronounced (**Fig. 1B** and **Fig. S2C**). This protein expression pattern of pathogenic *ADA2* variants was confirmed in ADA2^-/-^ U-937 cells transduced with the variants p.G47R, p.G47V and p.R169Q (**Fig. 1C** and **Fig. S2D**). In order to verify our finding in a larger number of *ADA2* variants, we analyzed the protein expression of mutant ADA2 by transfection of HEK293T cells. Overexpression of tagged WT ADA2 and several confirmed pathogenic variants uniformly led to predominant intracellular expression of HMW-ADA2 (63 kD) (**Fig. 1D**). In line with our findings in DADA2 HMDM, intracellular LMW-ADA2 was absent after overexpression of mutant ADA2 in HEK293T cells (**Fig. 1D**). Interestingly, this band was still present – albeit at lower intensity – in the variants p.N127Q and p.W326R, two variants not reported in DADA2 patients but created for the study of ADA2 glycosylation and dimerization, respectively (*11, 12*). The molecular weight of secreted wild-type (WT) ADA2 in the supernatant of HC HMDM or transfected HEK293T cells corresponded to the intracellular HMW form – the only form expressed in DADA2 HMDM (**Fig. 1B+D**). We confirmed the secretion of HMW-ADA2 in the monocytic cell lines U-937 and THP-1 (**Fig. S2E**). HMDM from DADA2 patient P8 showed residual secretion of HMW-ADA2 with a molecular weight equivalent to secreted WT ADA2 (**Fig. S2F**). This HMW-form of ADA2 also corresponded to the ADA2 form detected in human serum (**Fig. 1E**). Collectively, we show that HC HMDM intracellularly express a form of ADA2 that is lower in molecular weight than the secreted protein. This LMW-form is absent in DADA2 HMDM which only express the HMW-form intracellularly.

**Fig. 1:**
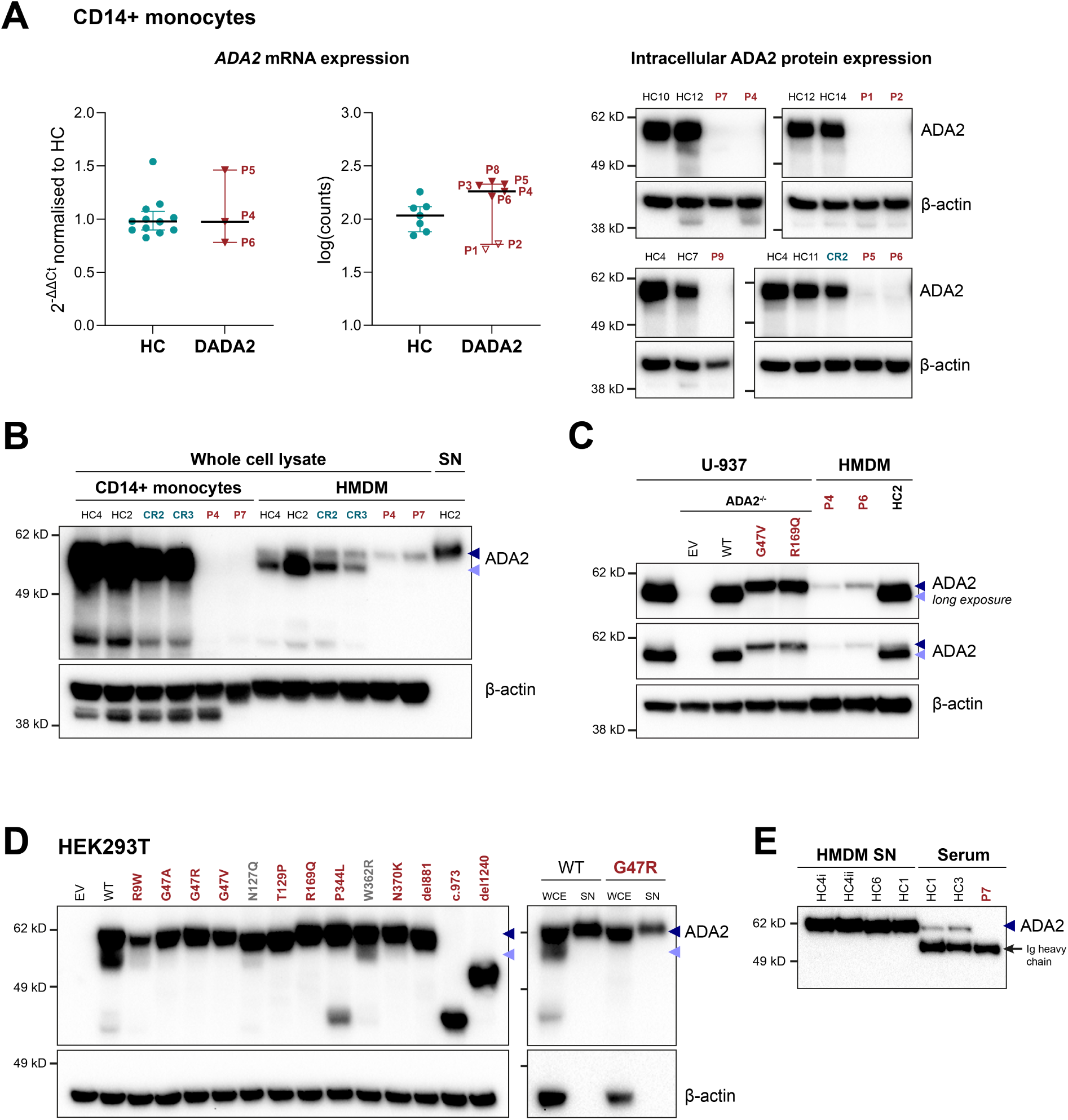
Expression of pathogenic ADA2 protein variants. **(A)** mRNA and intracellular protein expression of *ADA2* in CD14^+^ monocytes from healthy controls (HC) and DADA2 patients (P) by qPCR or bulk RNA-sequencing and western blot of whole cell lysates, respectively. The left plot shows mRNA expression determined by qPCR. *ADA2* mRNA expression was normalized to *GAPDH* and is depicted relative to the mean of HC samples. The right plot shows counts determined by RNA-sequencing. Dot plots show median and interquartile range. **(B)** ADA2 protein expression in whole cell lysates and supernatant (SN) of CD14^+^ monocytes and GM-CSF differentiated human monocyte-derived macrophages (HMDM) from HC, carriers (CR) and DADA2 patients (P) was determined by western blot. **(C)** ADA2 protein expression in whole cell lysates of transduced U-937 cells and HMDM from HC and DADA2 patients (P) was determined by western blot. **(D)** ADA2 protein expression in whole cell lysates of HEK293T cells after transfection with wild-type ADA2 and different pathogenic variants (*left*) and comparing whole cell lysates (WCE) and supernatant (SN) by western blot (*right*). **(E)** ADA2 protein expression in supernatants (SN) from healthy control human monocyte-derived macrophages (HMDM) and serum samples from healthy controls (HC) and DADA2 patients (P) by western blot using VeriBlot for detection. ADA2 detection using anti-ADA2 antibody clone EPR25430-131 (#ab288296; abcam) in all depicted western blots. Triangles indicate HMW-ADA2 (dark blue) and LMW-ADA2 (light blue). Legend: Ig, immunoglobulin; WT, wild type.

**Table 1:** Cohort characteristics. Genotype and clinical phenotype of the included DADA2 patients (P) and heterozygous carriers (CR).

| ID | Age<br>[years] | Sex | Genotype | Predominant phenotype |
| --- | --- | --- | --- | --- |
| P1 | 12 | M | c.973-2A>G/c.del1240-1442;<br>splice site (intron 6)/del exon 9 | Vasculitis, hepatosplenomegaly,<br>hypogammaglobulinemia |
| P2 | 14 | F | c.973-2A>G/c.del1240-1442;<br>splice site (intron 6)/del exon 9 | Vasculitis, warts, arthralgia,<br>hypo-gammaglobulinemia, |
| P3 | 26 | M | c.140G>T/c.del1240-1442;<br>p.G47V/del exon 9 | Bone marrow failure, hypogam-<br>maglobulinemia,<br>hepatosplenomegaly |
| P4 | 26 | M | c.140G>T/c.140G>T;<br>p.G47V/p.G47V | Hypogammaglobulinemia,<br>hepatosplenomegaly |
| P5 | 6 | F | c.140G>T/c.506G>A;<br>p.G47V/p.R169Q | Hypogammaglobulinemia,<br>hepatosplenomegaly, stroke |
| P6 | 9 | F | c.140G>T/c.506G>A;<br>p.G47V/p.R169Q | Bone marrow failure, hypogam-<br>maglobulinemia,<br>hepatosplenomegaly |
| P7 | 32 | F | c.973-2A>G/c.506G>A; splice<br>site (intron 6)/p.R169Q | Stroke, hypertension |
| P8 | 16 | M | c.139G>A/c.139G>A;<br>p.G47R/p.G47R | Stroke, vasculitis |
| P9 | 10 | F | c.973-2A>G/c.973-2A>G;<br>splice site (intron 6)/splice site<br>(intron 6) | Vasculitis,<br>hypogammaglobulinemia |
| P10 | 50 | F | c.973-2A>G/c.506G>A; splice<br>site (intron 6)/p.R169Q | Vasculitis, leukopenia |
| CR1 | 39 | F | WT/c. del1240-1442; WT/del<br>exon 9 |  |
| CR2 | 38 | F | WT/c.140G>T; WT/p.G47V |  |
| CR3 | 42 | M | WT/c.506G>A; WT/p.R169Q |  |

### Human monocyte-derived macrophages intracellularly express a hypoglycosylated low-molecular-weight form of ADA2 that is absent in DADA2 macrophages

Differences in the molecular weight of a protein can be caused by posttranslational protein modifications and N-glycosylation of ADA2 has previously been confirmed (*11, 13, 18*). Therefore, we initially hypothesized that the LMW-form of ADA2 unique to cells expressing WT ADA2 corresponded to glycan-free ADA2. To test this, we performed glycan removal on lysates and supernatants containing ADA2 protein (**Fig. 2A**). Complete removal of N-glycans by Peptide-N-Glycosidase F (PNGase F) yielded a difference in molecular weight of 7 kD between HMW-ADA2 and glycan-free ADA2 both overexpressed in HEK293T cells and endogenously expressed in the monocytic cell lines U-937 and THP-1 as well as HMDM (**Fig. 2B+C**), revealing that LMW-ADA2 was in fact slightly larger than glycan-free ADA2. Importantly, the anti-ADA2 antibody clone EPR25430-131 (#ab288296; abcam) was superior in detecting LMW-ADA2 but poorly detected glycan-free ADA2 (**Fig. S3A**). Glycan removal by PNGase F eliminated the differences in molecular weight between the LMW-form and the HMW-form in HMDM, yielding a single band of glycan-free ADA2 at 53 kD (**Fig. 2C-E**). Hence, the difference in molecular weight between intracellular WT ADA2 (LMW) and secreted WT ADA2 or intracellular pathogenic ADA2 (both HMW) was due to differential protein glycosylation. Of note, glycan removal yielded a smaller band of 53 kD in addition to the glycan-removed 56 kD protein for tagged WT ADA2 overexpressed in HEK293T cells that was absent for all tested pathogenic variants and did not appear after inhibition of N-glycosylation by tunicamycin (**Fig. 2B**). As this band was absent after glycan removal of endogenous WT ADA2, its physiological relevance is unclear. Overall, we showed that the difference in size between HMW- and LMW-ADA2 is due to differential protein glycosylation.

**Fig. 2:**
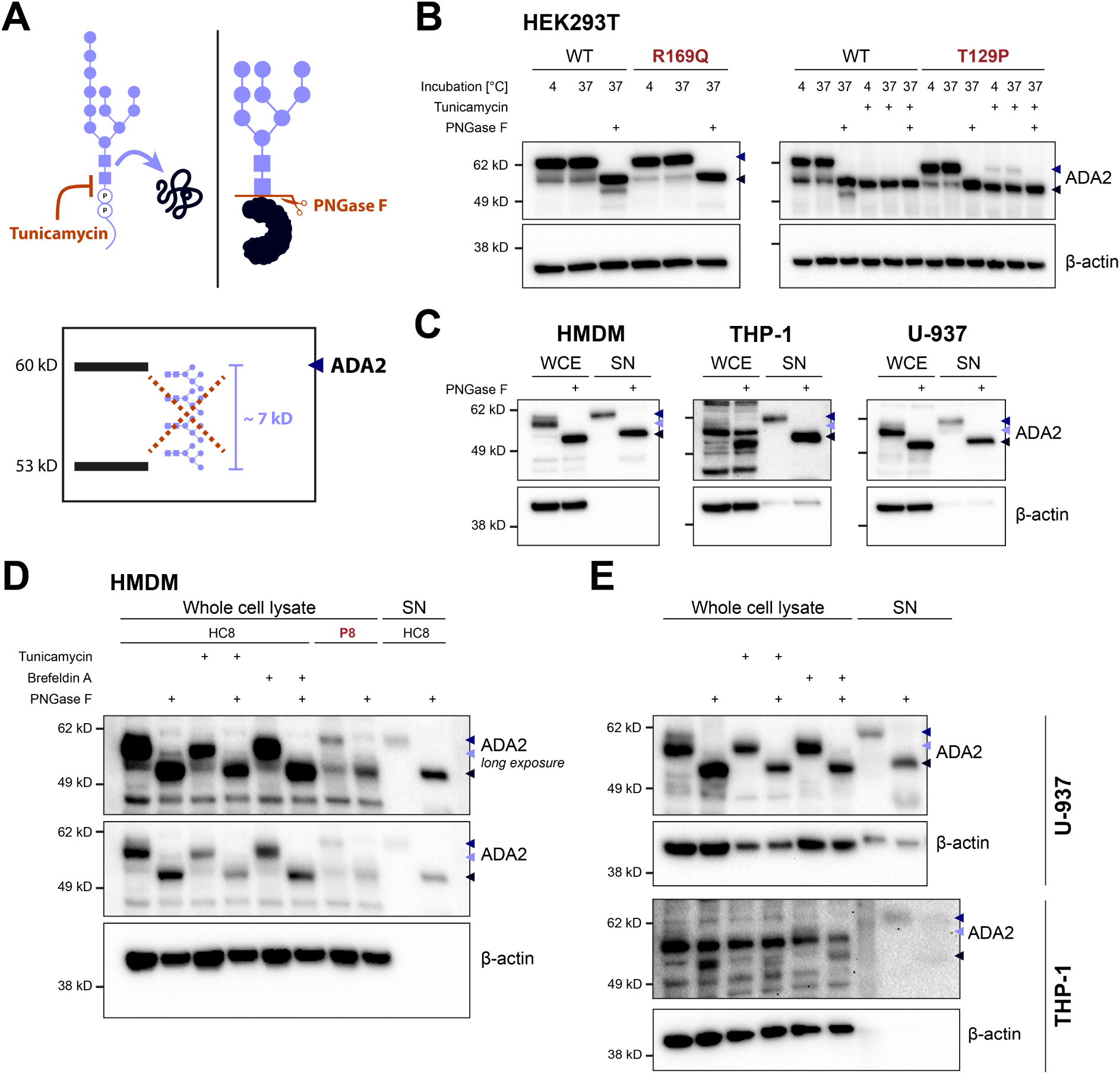
N-glycosylation of ADA2. **(A)** Schematic illustration of inhibition of glycosylation by tunicamycin and glycan removal by PNGase F. **(B)** Western blot of wild-type (WT) ADA2 and the pathogenic variants p.R169Q and p.T129P transfected into HEK293T cells after incubation with or without 2.5 µg/mL tunicamycin for 24 h and/or glycan removal by PNGase F. **(C)** ADA2 protein expression by western blot in whole cell lysates (WCE) and supernatants (SN) of human monocyte-derived macrophages (HMDM), U-937 cells and THP-1 cells. **(D)** ADA2 protein expression in whole cell lysates and supernatants (SN) from GM-CSF-differentiated HMDM of healthy control HC8 and DADA2 patient P8 left untreated or treated with 5 µg/mL tunicamycin or 1 µg/mL brefeldin A for 24 h. **(E)** ADA2 protein expression in whole cell lysates and supernatants (SN) from U-937 and THP-1 cells left untreated or treated with 2.5 µg/mL tunicamycin or 1 µg/mL brefeldin A for 24 h. Glycan removal was performed by incubation with PNGase F for 1 hour at 37°C under denaturing conditions in all experiments. ADA2 detection using anti-ADA2 (#HPA007888, Sigma-Aldrich) in all depicted western blots. Triangles indicate HMW-ADA2 (dark blue), LMW-ADA2 (light blue) and glycan-free ADA2 (black).

### N-glycosylated ADA2 undergoes glycan trimming in the ER as part of protein quality control

In order to better characterize the glycosylation of endogenous ADA2, we inhibited N-glycosylation in HMDM and the monocytic cell lines U-937 and THP-1 by 24-hour treatment with tunicamycin. We found that this treatment only acted on the fainter HMW-band while LMW-ADA2 remained unchanged in size (**Fig. 2D+E**). In accordance with previous findings (*13*), we showed that adenosine deaminase activity of ADA2 was preserved after glycan removal while treatment with tunicamycin reduced enzymatic activity. This was true for ADA2 overexpressed in HEK293T cells as well as endogenous ADA2 from HC HMDM (**Fig. S3B+C**). These findings indicated that N-glycosylation was required for adequate folding of the ADA2 protein. The ER glucosidases I and II form part of the ER quality control by removing glycosyl residues from accurately folded proteins prior to transfer to the Golgi apparatus (*20*). Treatment with castanospermine, an inhibitor of the ER glucosidases I and II, caused an additional increase in the size of HMW-ADA2 in both HC and DADA2 HMDM (**Fig. 3A**), confirming trimming of N-glycosylated ADA2 in the ER. Since the same shift in molecular weight was observed in DADA2 HMDM, we concluded that processing of WT and mutant does not yet diverge at this early stage of the secretory pathway. As seen previously upon incubation with tunicamycin, the LMW-form of ADA2, which is absent in DADA2 HMDM, was unaffected by this treatment (**Fig. 3A**). We confirmed this finding in the three ADA2-expressing cell lines U-937, THP-1 and Jurkat (**Fig. 3B**). Since hypoglycosylated LMW-ADA2 was still detected after 24-hour treatment with tunicamycin – an inhibitor of the initial step of N-glycosylation – we concluded that glycosylation and trimming of the residual LMW-form must have occurred prior to the initiation of the incubation period, suggesting high stability of LMW-ADA2. Accordingly, LMW-ADA2 was responsive to prolonged incubation with tunicamycin at a lower concentration (**Fig. 3B**). The same was true for inhibition of ER glucosidases: the more stable LMW-ADA2 was unaffected by 24-hour incubation with castanospermine and the detected LMW-form had likely already matured before the beginning of the incubation period. We confirmed the increased stability of LMW-ADA2 compared to HMW-ADA2 by cycloheximide chase assay (**Fig. 3C** and **Fig. S3D**). Besides, this experiment revealed that HMW-ADA2 protein expressed by HEK293T cells transfected with pathogenic *ADA2* variants showed increased intracellular stability compared with the WT protein. This was also true after inhibition of N-glycosylation and after an increased incubation period of 48 hours (**Fig. 3C** and **Fig. S3E**). Since ADA2 is a secreted protein, protein secretion must be taken into account when assessing the intracellular signal reduction in this assay over time (**Fig. S3F**). Impaired secretion has previously been established as a characteristic of pathogenic ADA2 variants (*2, 14*). We confirmed the secretory defect in HMDM and serum from our DADA2 cohort (**Fig. 3D**). Overexpression in HEK293T cells recapitulated the endogenous behavior of the respective variants well (**Fig. 3D**). Comparing the levels of increased ADA2 stability observed by cycloheximide chase assay with residual protein secretion, the variants p.T129P and p.R169Q that exhibited complete absence of secretion did indeed show the highest levels of intracellular ADA2 24 hours after inhibition of protein synthesis (**Fig. 3C** and **Fig. S3D**). However, intracellular levels of the secreted mutant p.P344L were still higher compared with WT.

**Fig. 3:**
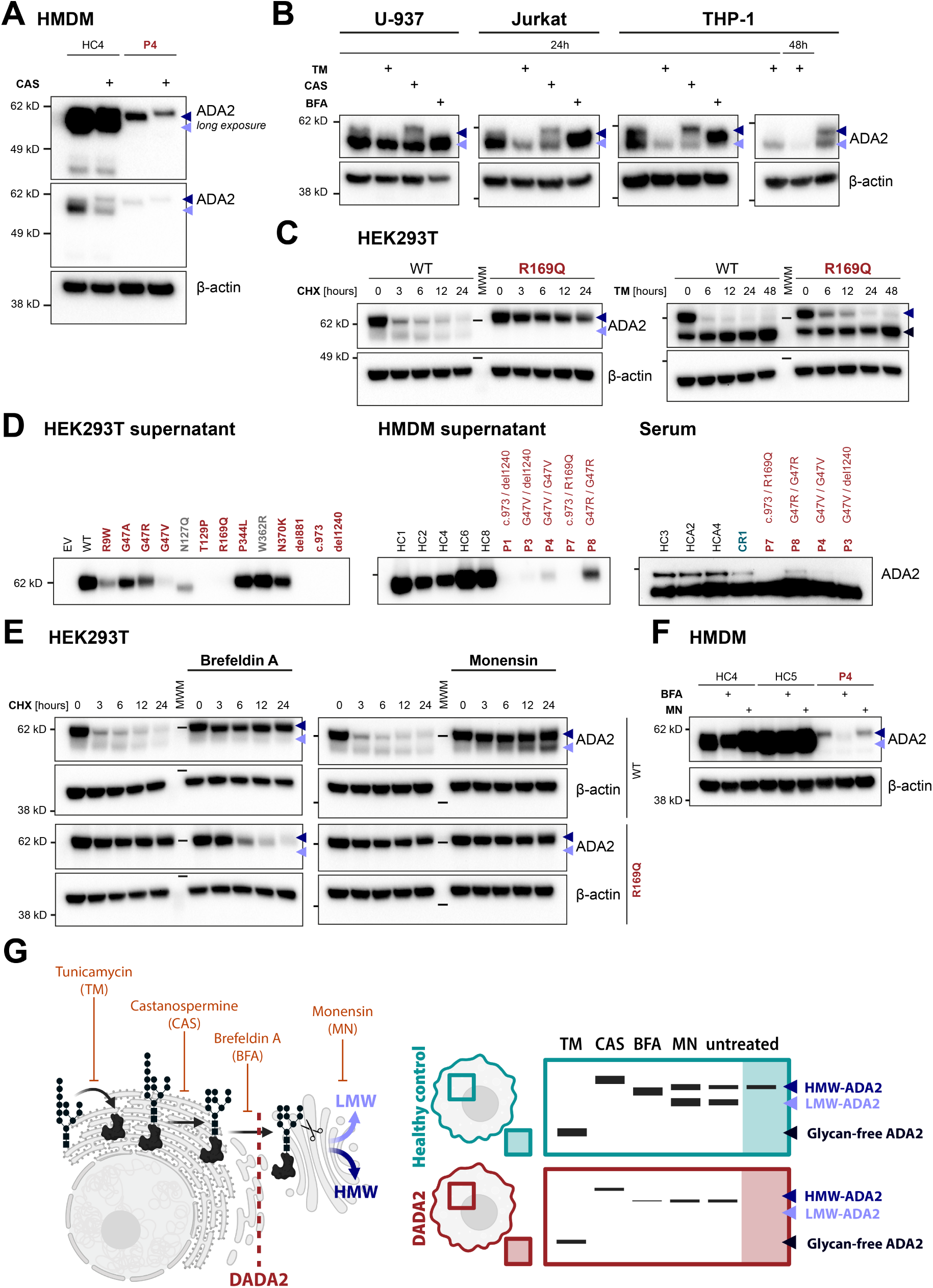
Glycan processing of ADA2 in the ER. **(A)** ADA2 protein expression by western blot in GM-CSF-differentiated HMDM from healthy control HC4 and DADA2 patient P4 after incubation with 100 µg/mL castanospermine for 24 h. **(B)** ADA2 protein expression upon inhibition of N-glycosylation, ER glucosidases I and II, or ER to Golgi trafficking. U-937, THP-1 and Jurkat cells were incubated with 2.5 µg/mL tunicamycin (TM), 100 µg/mL castanospermine (CAS) or 1 µg/mL brefeldin A (BFA) for 24 h or 48 h. **(C)** HEK293T cells were transfected with wild-type (WT) ADA2 or the pathogenic variant p.R169Q and treated with 500 µg/mL cycloheximide (CHX) or 2.5 µg/mL tunicamycin (TM) over 24-48 h. Whole cell lysates were generated at the indicated time points and immunoblotted for ADA2 expression. **(D)** ADA2 protein levels were determined in supernatants of M-CSF-differentiated human monocyte-derived macrophages (HMDM) from healthy controls (HC) or DADA2 patients (P) or HEK293T cells transfected with wild-type (WT) *ADA2* or different pathogenic *ADA2* variants. The patients’ genotypes are indicated above. Western blot of serum samples from healthy controls (HC), heterozygous DADA2 carriers (CR) and DADA2 patients (P) detected by anti-ADA2 antibody clone EPR25430-131 (#ab288296; abcam) and VeriBlot. **(E)** HEK293T cells were transfected with wild-type (WT) ADA2 or different pathogenic variants and treated with 500 µg/mL cycloheximide (CHX) ± 1 µg/mL brefeldin A or 2 µg/mL monensin over 24 h. Whole cell lysates were generated at the indicated time points and immunoblotted for ADA2 expression. **(F)** ADA2 protein expression in GM-CSF-differentiated human monocyte-derived macrophages from healthy controls (HC) or DADA2 patient P4 left untreated or treated with 1 µg/mL brefeldin A (BFA) or 2 µg/mL monensin (MN). **(G)** Schematic overview of trafficking and processing of ADA2 in healthy control and DADA2 cells. ADA2 detection using anti-ADA2 antibody clone EPR25430-131 (#ab288296; abcam) in panels A, B, C (*left*), D, E and F and using anti-ADA2 (#HPA007888, Sigma-Aldrich) in panel C (*right*). Triangles indicate HMW-ADA2 (dark blue), LMW-ADA2 (light blue) and glycan-free ADA2 (black). Legend: BFA, brefeldin A, CAS, castanospermine, EV, empty vector; MN, monensin, MWM, molecular weight marker; TM, tunicamycin; WT, wild type.

We next blocked the secretory pathway to impair secretion of WT ADA2 and thereby confirmed that the secretory defect strongly contributed to the increased intracellular protein levels of mutant ADA2 (**Fig. 3E** and **Fig. S4**). While the secretory defect of mutant ADA2 was phenocopied by WT ADA2 upon incubation with brefeldin A (BFA), we also found that this treatment caused increased degradation of the mutants p.G47V, p.T129P and p.R169Q (**Fig. 3E** and **Fig. S4**). BFA inhibits ER to Golgi trafficking. We observed that treatment of WT ADA2-expressing cells with BFA yielded a single ADA2 band with a molecular weight between that of HMW- and LMW-ADA2 (**Fig. 3B+E**), suggesting that both HMW- and LMW-ADA2 are the result of further glycan processing in the Golgi. Indeed, inhibition of trans Golgi transport by monensin (*21, 22*) did not impair the generation of LMW-ADA2 in cells overexpressing WT ADA2 (**Fig. 3E**). The differential behavior in response to the secretory pathway inhibitors BFA and monensin was confirmed in DADA2 HMDM (**Fig. 3F**). The absence of LMW-ADA2 upon inhibition of ER to Golgi trafficking and the increase in LMW-ADA2 upon inhibition of trans Golgi transport supported our hypothesis that LMW- and HMW-ADA2 are generated from a common precursor glycoform after transport to the Golgi apparatus (**Fig. 3G**).

In summary, we showed that N-glycosylation is required for folding and secretion of ADA2 and that both WT and mutant ADA2 are subject to processing by the ER glucosidases I and II while only WT ADA2 undergoes further glycan trimming in the Golgi apparatus.

### The pathogenic ADA2 variant p.R169Q shows increased binding to proteins of the protein folding machinery

After verifying the expression of LMW- and HMW-ADA2 in cells expressing WT ADA2 and an impairment of glycan-processing in DADA2 cells, our aim was to further characterize the different glycoforms of ADA2 in the second part of this study: (i) retained mutant HMW-ADA2, (ii) secreted WT HMW-ADA2 and (iii) intracellular WT LMW-ADA2.

First, we performed immunoprecipitation followed by mass spectrometry (IP-MS) to analyze the interactome of WT ADA2 and the pathogenic ADA2 variant p.R169Q transfected into HEK293T cells. p.R169Q displayed increased interaction with proteins involved in the ER quality control machinery for glycoproteins (**Fig. 4A** and **Table S1**). Upon inhibition of N-glycosylation by tunicamycin, we observed a similar increase also for the WT protein (**Fig. 4A** and **Table S1**), highlighting the importance of adequate N-glycosylation for successful folding and secretion of ADA2. Additionally, we observed an increase in lysosomal proteins in the ADA2 interactome after treatment with tunicamycin, suggesting autophagic degradation of non-glycosylated ADA2. Among the induced interactors were several proteins of the chaperone-mediated autophagy pathway, e.g. LAMP2 and HSPA13, and we verified the presence of a KFERQ-related motif (QKFVE, amino acids 274-278) in the sequence of ADA2 (*23*). IP-MS of HC and DADA2 HMDM did not yield any specific binders, which was possibly due to low protein abundance in the pull down of endogenous ADA2 (**Fig. S5A**). Notably, IP preferentially pulled down the HMW-form of ADA2 (**Fig. S5B+C**). The experiment consequently did not allow us to draw conclusions about the interactome and potential distinctive function of LMW-ADA2.

**Fig. 4:**
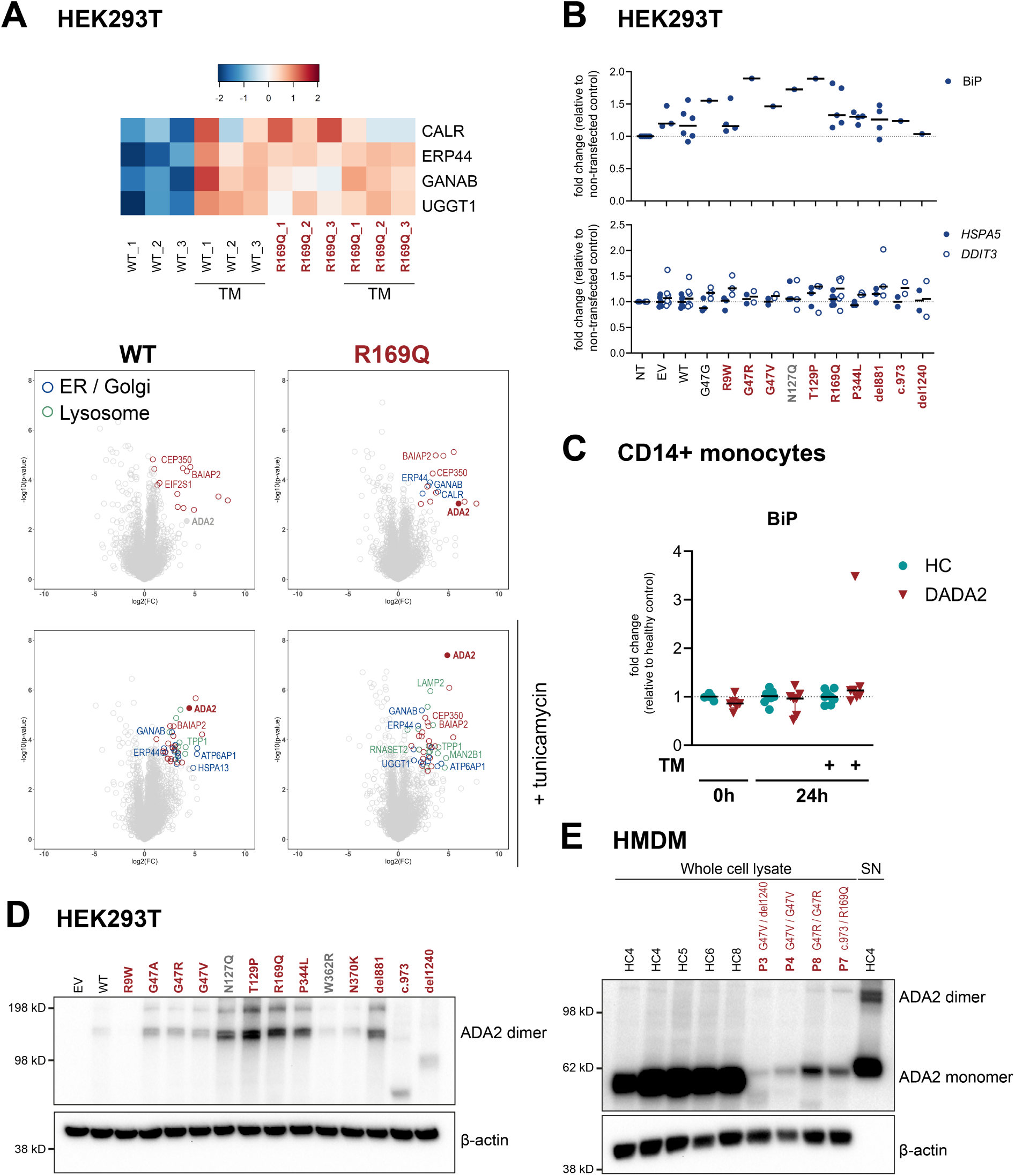
ER stress in DADA2. **(A)** Interactome analysis of wild-type (WT) ADA2 and the pathogenic variant p.R169Q overexpressed in HEK293T cells treated with or without 2.5 µg/mL tunicamycin (TM) for 24 h was performed by immunoprecipitation followed by mass spectrometry. Heatmap shows z-score of protein abundance. Binders significantly pulled down compared to isotype control with a false discovery rate of 5% are highlighted by colored circles. Localization of the interacting proteins to the ER/Golgi (blue) or lysosome (green) is indicated by color. **(B)** ER stress response in HEK293T cells transfected with wild-type (WT) *ADA2* and different pathogenic variants. Protein expression of BiP (*top*) and mRNA expression of *HSPA5* and *DDIT3* (*bottom*) were determined 48h after transfection. A representative western blot is shown in **Fig. S5F**. mRNA expression was normalized to *GAPDH* and is depicted as 2^-ΔΔCt^ relative to non-transfected control samples. Results from 6 independent experiments are shown. Dot plots show median. **(C)** ER stress response was evaluated by protein expression of BiP in whole cell lysates of CD14^+^ monocytes at baseline or after 24 h incubation with or without 1 µg/mL tunicamycin (TM). BiP expression was normalized to β-actin. Dot plots show median and interquartile range. The corresponding western blots are shown in **Fig. S6**. **(D)** Formation of intracellular ADA2 dimers in HEK293T cells transfected with wild-type (WT) ADA2 and different pathogenic *ADA2* variants. The blot is equivalent to the one shown in Fig. 1D. **(E)** ADA2 protein expression and intracellular dimer formation in GM-CSF differentiated human monocyte-derived macrophages from healthy controls (HC) and DADA2 patients (P). ADA2 detection using anti-ADA2 antibody clone EPR25430-131 (#ab288296; abcam) in all depicted western blots. Legend: EV, empty vector; FC, fold change; NT, non-transfected; WT, wild type.

Given the increased binding of mutant ADA2 to the ER folding machinery, we verified whether pathogenic *ADA2* variants induced an increased ER stress response. In line with previous reports (*14*), we observed that some pathogenic *ADA2* variants caused signs of activation of an ER stress response, measured by increased levels of CHOP (*DDIT3*) and BiP (*HSPA5*) (**Fig. S5D**). These differences were however no longer present after down-titrating the amount of transfected DNA to achieve near-physiological protein levels (**Fig. 4B** and **Fig. S5E+F**) and neither was there a difference in BiP expression in CD14^+^ monocytes from DADA2 patients compared to healthy controls (**Fig. 4C** and **Fig. S6**). In line with other studies (*8, 24*), we showed that protein misfolding led to intracellular aggregate formation of mutant ADA2 (**Figure 4D+E**). In summary, our experiments revealed the interaction of the ADA2 variant p.R169Q overexpressed in HEK293T cells with mediators of the folding machinery and ER quality control despite lack of increased ER stress in the presence of pathogenic ADA2 variants. The interactome analyses were inconclusive regarding the characterization of LMW-ADA2.

### Wild-type HMW-ADA2 is subject to additional glycan maturation prior to secretion

Next, we aimed to further explore the glycan structure of HMW-ADA2. We hypothesized that the ER retention of most pathogenic protein variants of ADA2 leads to incomplete glycan processing. To examine the glycosylation pattern of WT and mutant ADA2, we performed additional glycan removal experiments in ADA2^-/-^ U-937 cells transduced pathogenic *ADA2* variants. We chose a monocytic cell line for these experiments to minimize errors due to differential expression of glycosylation enzymes in different cell types.

First, we confirmed that complete N-glycan removal by PNGase F cancelled out the differences in molecular weight between HMW- and LMW-ADA2, confirming that they are due to N-glycosylation (**Fig. 5A**). Endo H is another endoglycosidase that cleaves high mannose glycans and – unlike PNGase F – does not remove complex glycans. We found that WT and mutant ADA2 were both sensitive to glycan removal by Endo H (**Fig. 5A+B**), hinting at the presence of high mannose oligosaccharides in both LMW- and HMW-ADA2 glycans. In contrast with PNGase F treatment, glycan removal by Endo H revealed differences between WT and mutant ADA2: Compared with mutant ADA2, Endo H-treated WT ADA2 visualized as a smear on western blot as opposed to the clean single band observed for mutant ADA2 in U-937 cells and HMDM (**Fig. 5A+B**). This suggests that WT ADA2 undergoes additional glycan processing as complex glycans are less sensitive to Endo H. Since the size of LMW-ADA2 makes the presence of additional carbohydrate residues unlikely, we concluded that in WT ADA2-expressing cells, the glycan trees of HMW-ADA2 mature into complex glycans, generating HMW(c)-ADA2, the HMW-form observed in HC HMDM (**Fig. 5C**). For all tested pathogenic variants, glycan removal by Endo H yielded a clean band very close in molecular weight to complete removal of N-glycans by PNGase F (**Fig. 5A**). Consequently, mutant HMW-ADA2 is likely not subject to any glycan maturation beyond the ER, which is in agreement with the postulated ER retention of pathogenic ADA2 protein variants. This finding also implies that WT HMW(c)-ADA2 and mutant HMW-ADA2 – while very similar in molecular weight –differ in their glycan structures and do not represent the same ADA2 glycoform.

**Fig. 5:**
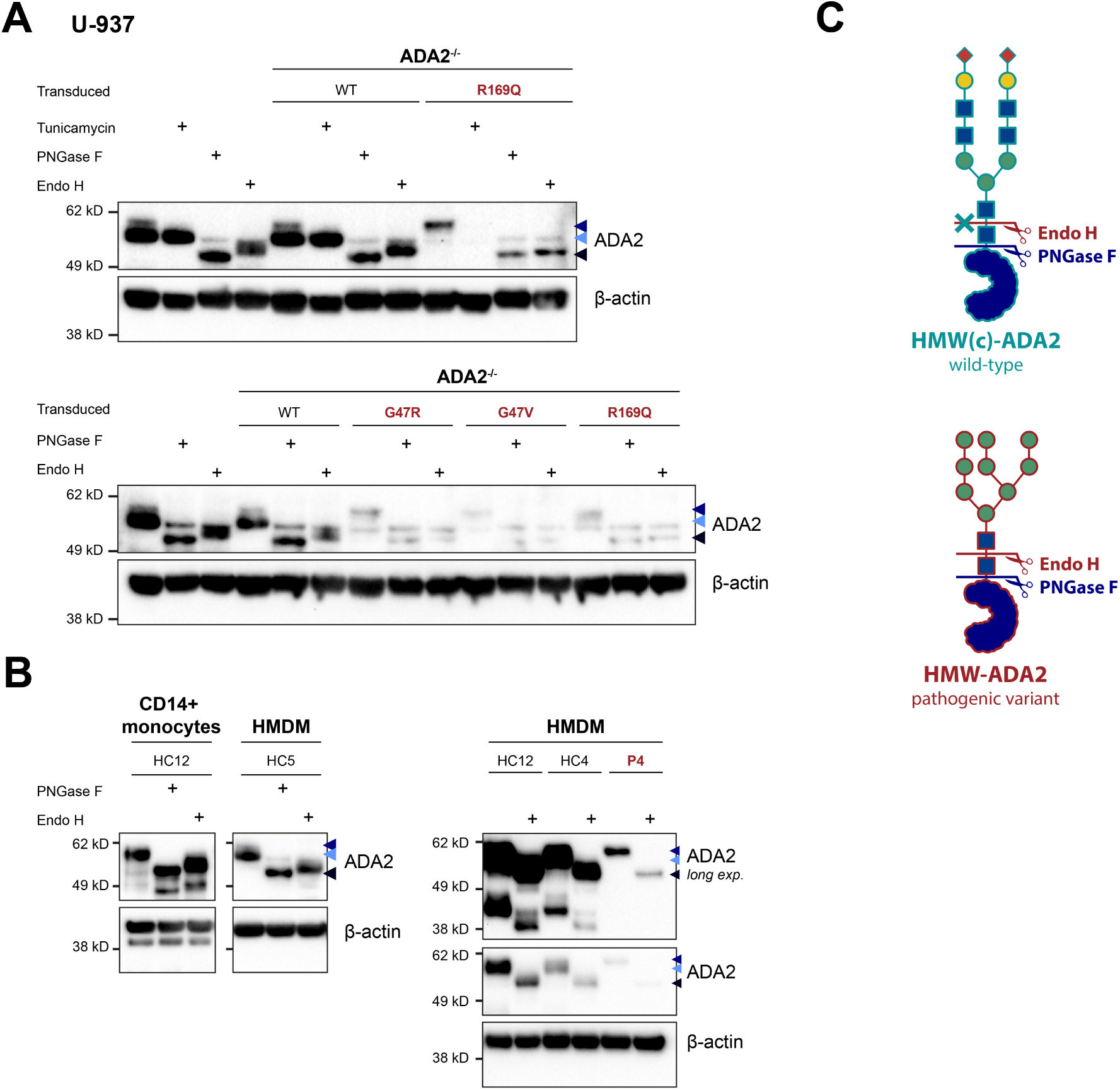
Glycan processing of HMW(c)-ADA2. ADA2 protein expression by western blot after glycan removal in ADA2^WT/WT^ and ADA2^-/-^ U-937 cells transduced with pathogenic *ADA2* variants **(A)** and CD14^+^ monocytes or GM-CSF-differentiated HMDM from healthy controls (HC) and DADA2 patient P4 **(B)** by incubation with PNGase F or Endo H for 1 hour at 37 °C under denaturing conditions. **(C)** Cutting sites of PNGase F and Endo H. Endo H does not cleave complex glycans (*top*). Carbohydrate residues are depicted according to the Symbol Nomenclature for Glycans (SNFG) (*20*). ADA2 detection using anti-ADA2 antibody clone EPR25430-131 (#ab288296; abcam) in panel B (*right*) and using anti-ADA2 (#HPA007888, Sigma-Aldrich) in panels A and B (*left*). Triangles indicate HMW-ADA2 (dark blue), LMW-ADA2 (light blue) and glycan-free ADA2 (black).

Collectively, our findings suggest that the N-glycan structures of ADA2 contain high-mannose oligosaccharides and that WT HMW(c)-ADA2 contains complex glycan branches while mutant HMW-ADA2 does not undergo advanced glycan maturation.

### LMW-ADA2 is generated by glycan trimming through alpha-mannosidases

The addition of complex glycan structures is usually preceded by the removal of α-linked mannose residues in the late ER and Golgi by a set of α-mannosidases (*25*). We therefore hypothesized that mutant ADA2 is inaccessible to glycan-trimming by endogenous α-mannosidases, accounting for the absence of complex glycan structures as well as the absence of LMW-ADA2. To test this hypothesis, we incubated WT and mutant ADA2-expressing U-937 cells with different inhibitors of glycan-processing enzymes. The corresponding western blots were performed in a capillary-based separation and detection system to allow for a more precise distinction of the different molecular weight peaks. After synthesis and N-glycan transfer in the ER, proteins are initially trimmed by the ER α-glucosidases I and II that are inhibited by castanospermine (*20*). As shown in HMDM, castanospermine caused an increase in the molecular weight of HMW-ADA2 (**Fig. S7**). By increasing the incubation time ≥ 48 hours we observed a shift also for LMW-ADA2, confirming the superior stability of LMW-ADA2. The change in molecular weight of both HMW- and LMW-ADA2 upon inhibition of the ER α-glucosidases I and II supported our hypothesis that the processing of the two glycoforms only diverges after exiting the ER. After passing the quality control mechanisms in the ER, proteins can be trimmed further by different α-mannosidases in the ER and Golgi. Kifunensine inhibits class I α-mannosidases that are typically found in the late ER and cis-Golgi and cleave α-1,2-linked mannose residues (*20*). Incubation of U-937 cells with kifunensine led to an increase in the molecular weight of LMW-ADA2 (**Fig. 6A** and **Fig. S7**). After 96 hours, LMW-ADA2 almost overlay HMW-ADA2 (**Fig. 6A**). Concomitant incubation with swainsonine led to similar results. However, incubation with swainsonine alone also increased the molecular weight of LMW-ADA2, suggesting that multiple mannosidases are involved in the trimming of LMW-ADA2. Swainsonine inhibits both α-mannosidase II in the medial Golgi and the lysosomal α-mannosidase (*26*). To better understand whether also the lysosomal enzyme is involved in the trimming of LMW-ADA2, we analyzed HMDM from a patient with α-mannosidosis, harbouring biallelic mutations in *MAN2B1* encoding the lysosomal α-mannosidase (*27*). We found that the patient’s cells phenocopied HC cells treated with the class II α-mannosidase inhibitor swainsonine and that swainsonine treatment did not have an additional effect on the molecular weight of LMW-ADA2 expressed in the lysosomal α-mannosidase-deficient cells (**Fig. 6B**). This finding suggests that LMW-ADA2 undergoes processing not only in the ER and Golgi but also in the lysosomes. Finally, as a proof of principle, we performed glycan removal by α1-2,3,6 mannosidase on WT ADA2. This experiment yielded a single band slightly lower but very close in molecular weight to LMW-ADA2 (**Fig. 6C**). This is in line with the findings of the previous experiments since the inhibited glycan-processing enzymes typically do not cleave α-1,6-linked mannose residues, which accounts for the lower molecular weight of α1-2,3,6 mannosidase-treated ADA2 compared with LMW-ADA2 (**Fig. 6D+E**). Importantly, the glycan structures of LMW-ADA2 predicted by this set of experiments corresponded with the glycan profile determined in another study (**Fig. S8A**) (*28*).

**Fig. 6:**
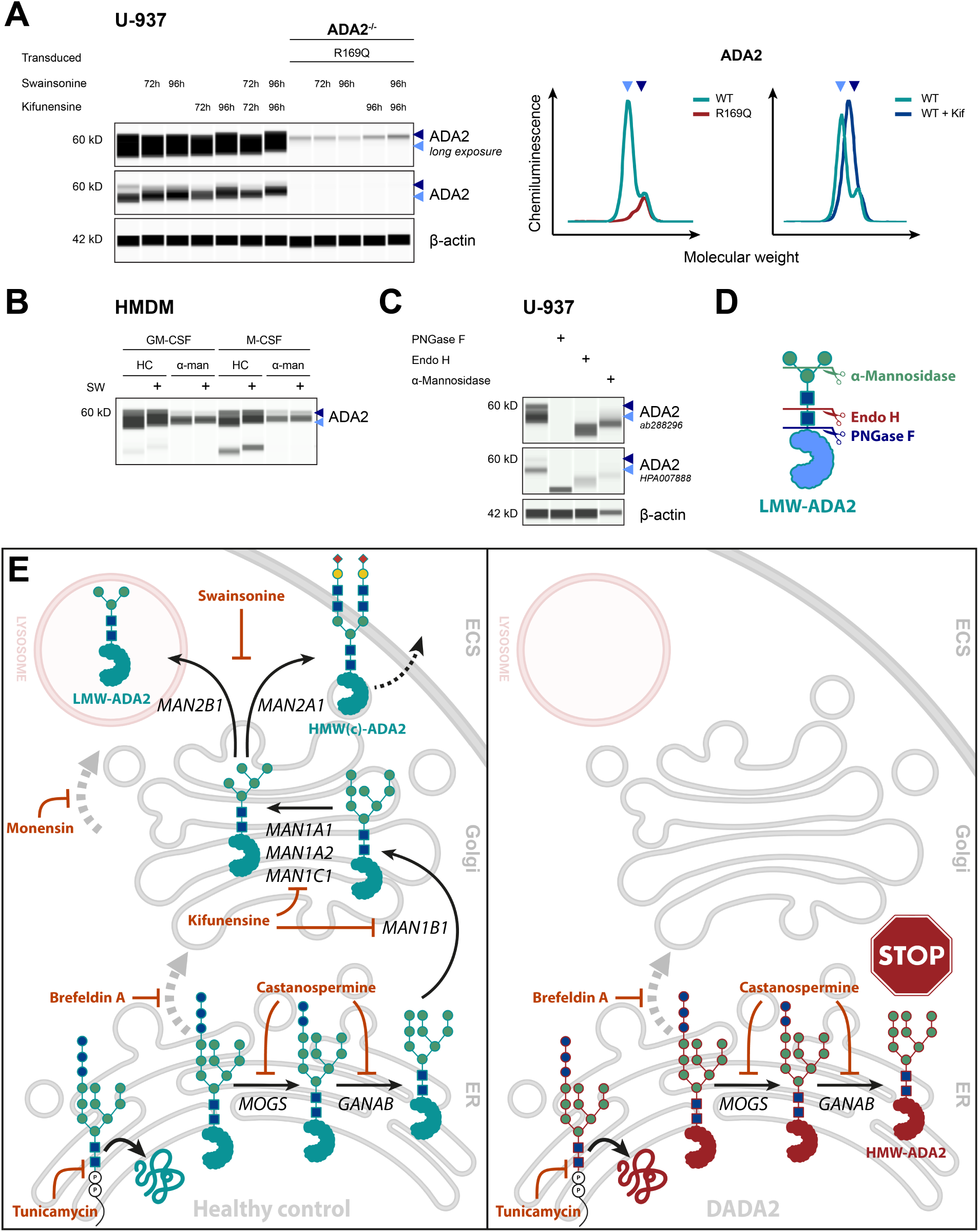
Glycan processing of LMW-ADA2. **(A)** α-mannosidases were inhibited by 72-96 h incubation of ADA2^WT/WT^ and ADA2^-/-^ U-937 cells transduced with R169Q with 375 nM kifunensine and/or 10 µM swainsonine. ADA2 protein expression was determined by western blot. The histograms indicate molecular weight peaks as determined by Simple Western. The original graphs are provided in **Fig. S7**. **(B)** ADA2 protein expression by western blot GM-CSF- and M-CSF-differentiated HMDM from a healthy control (HC) and a patient with α-mannosidosis (α-man) after incubation with 10 µM swainsonine (SW) for 96 h. **(C)** ADA2 protein expression by western blot after glycan removal in ADA2^WT/WT^ U-937 cells by incubation with PNGase F, Endo H or α1-2,3,6 Mannosidase for 1 hour at 37 °C. **(D)** Cleavage sites of PNGase F, Endo H and α-Mannosidase. **(E)** Schematic overview of the trafficking and glycan modifications of ADA2 expressed in healthy control (*left*) and DADA2 (*right*) human monocyte-derived macrophages based on the findings presented in this article. *MOGS* encodes ER α-glucosidase I, *GANAB* encodes ER α-glucosidase II, *MAN1B1* encodes ER α-1,2-mannosidase, *MAN1A1*, *MAN1A2* and *MAN1C1* encode Golgi α-1,2-mannosidases, *MAN2A1* encodes Golgi α-mannosidase II, and *MAN2B1* encodes lysosomal α-mannosidase. Mechanisms of action of the inhibitors used for the *in vitro* study of glycan processing are depicted in orange. Carbohydrate residues are depicted according to the Symbol Nomenclature for Glycans (SNFG) (*20*). ADA2 detection using anti-ADA2 antibody clone EPR25430-131 (#ab288296; abcam) in panels A, B and C (*top*) and using anti-ADA2 (#HPA007888, Sigma-Aldrich) in panel C (*bottom*). Triangles indicate HMW-ADA2 (dark blue), LMW-ADA2 (light blue) and glycan-free ADA2 (black). Legend: ECS, extracellular space; ER, endoplasmic reticulum.

In summary, we demonstrated that LMW-ADA2 is generated from HMW-ADA2 by glycan-processing through multiple α-mannosidases in the ER, Golgi and lysosomal compartments.

### LMW-ADA2 localizes to the lysosomes and can be generated from WT HMW(c)-ADA2 in ADA2-deficient cells

To further support the processing of LMW-ADA2 by the lysosomal α-mannosidase, we next aimed to verify the lysosomal localization of the LMW-glycoform in monocytic cells. To this end, we performed immunofluorescence microscopy of U-937 cells expressing WT ADA2 or the pathogenic ADA2 variant p.R169Q. In cells expressing the WT protein, ADA2 colocalized with the lysosomal marker LAMP1 (**Fig. 7A** and **Fig. S8B**). Moreover, the experiment confirmed ER retention of the mutant protein while WT ADA2 was found in the ER, Golgi and lysosomal compartments (**Fig. 7A**). To verify that the intracellular glycoform of ADA2 localizing to the lysosomes was indeed LMW-ADA2, we then performed subcellular fractionation experiments. We showed that the lysosomal fraction contained only LMW-ADA2 (**Fig. 7B**). LMW-ADA2 was also present in the cytosolic and membrane fractions. HMW-ADA2 localized exclusively to the membranous fraction that also contains ER and Golgi membrane proteins (**Fig. 7B**).

**Fig. 7:**
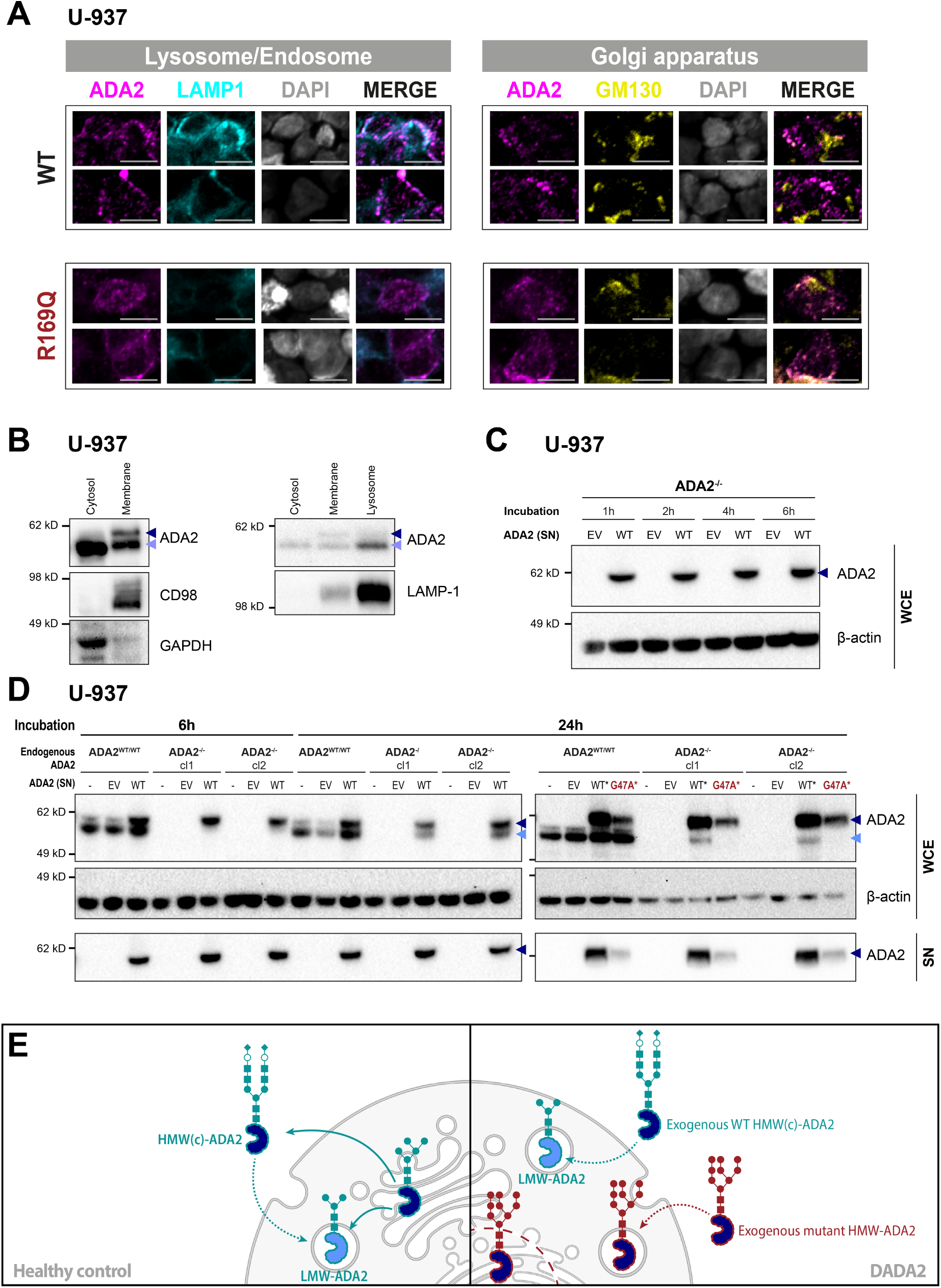
Localization and generation of LMW-ADA2. **(A)** Immunofluorescence microscopy of ADA2^WT/WT^ U-937 cells and ADA2^-/-^ U-937 transduced with the pathogenic variant p.R169Q stained for ADA2 and markers of lysosomal (LAMP1) and Golgi (GM130) compartments. **(B)** ADA2 protein expression in cytosolic, membranous and lysosomal fractions of U-937 cells by western blot. **(C)** Intracellular ADA2 uptake into ADA2^-/-^ U-937 cells after incubation with supernatants (SN) from HEK293T cells transfected with empty vector (EV) or wild-type (WT) *ADA2* for the indicated time. **(D)** Generation of intracellular LMW-ADA2 in ADA2^WT/WT^ or ADA2^-/-^ U-937 cells after incubation with supernatants (SN) from HEK293T cells transfected with empty vector (EV), wild-type (WT) *ADA2* or the pathogenic variant p.G47A for the indicated time. **(E)** Schematic illustration of uptake and processing of exogenous ADA2. ADA2 detection using anti-ADA2 antibody clone EPR25430-131 (#ab288296; abcam) in panels C and D and using anti-ADA2 (#HPA007888, Sigma-Aldrich) in panel B. Triangles indicate HMW-ADA2 (dark blue), LMW-ADA2 (light blue) and glycan-free ADA2 (black). Legend: cl, clone.

Besides intracellular transport, proteins can also enter the lysosomes via the endocytotic pathway. To evaluate whether this route allowed the uptake and processing of ADA2 from the extracellular space, we incubated ADA2^-/-^ U-937 cells with the supernatant of transfected HEK293T cells containing WT HMW(c)-ADA2. We first confirmed that exogenous HMW(c)-ADA2 protein was taken up by the ADA2-deficient cells (**Fig. 7C**). Next, we analyzed whether HMW(c)-ADA2 underwent intracellular glycan processing after uptake. Within the first 6 hours of incubation, HMW-ADA2 was the only glycoform found intracellularly (**Fig. 7D**). After 24 hours, we observed a strong signal for LMW-ADA2 in the whole cell lysate of the ADA2-deficient cells suggesting that exogenous HMW(c)-ADA2 was subject to intracellular glycan trimming after uptake (**Fig. 7D**). In accordance with our previous findings, LMW-ADA2 was not found in the supernatant of the incubated cells. This finding also mechanistically adds to our previous observation that LMW-ADA2 exhibits increased stability: Generation of LMW-ADA2 from internalized previously secreted extracellular HMW(c)-ADA2 represents an additional mechanism accounting for the prolonged presence of intracellular LMW-ADA2 after inhibition of protein synthesis or N-glycosylation.

Finally, we confirmed that the secreted pathogenic *ADA2* variant p.G47A was also taken up into the monocytic cells. Unlike the WT protein, the mutant protein was however not processed into LMW-ADA2 (**Fig. 7D**). These experiments confirm that LMW-ADA2 can be generated from exogenous extracellular ADA2 and that ADA2-deficient cells possess the glycan-processing machinery to generate LMW-ADA2 from the WT protein (**Fig. 7E**).

### Absence of LMW-ADA2 correlates with impaired ADA2 enzyme activity and predicts the pathogenicity of *ADA2* variants

After characterizing LMW-ADA2 as the intracellular glycoform of ADA2 and establishing its absence as a feature of DADA2 cells, we validated our findings on a larger set of *ADA2* variants. For this, we analyzed a total of 34 *ADA2* variants, including 18 variants found in homozygous state in DADA2 patients and two variants predicted to be benign with homozygous individuals documented in the gnomAD database (p.H335R and p. R230Q) (gnomAD v4.1.0, https://gnomad.broadinstitute.org). Our data set also included all the *ADA2* variants previously described as DADA2-associated despite an ADA2 enzyme activity above 25% of WT ADA2 (*29*). The included variants cover all the domains of the ADA2 protein (**Fig. 8A**). The variants were overexpressed in HEK293T cells and studied for LMW-ADA2 expression, ADA2 protein secretion and both extracellular and intracellular ADA2 enzyme activity (**Fig. 8B**). In line with our studies on the trafficking of LMW-ADA2, this glycoform was exclusively processed from variants that also showed residual ADA2 secretion (**Fig. 8B**). The greatest part of the pathogenic variants with residual secretion lacking enzymatic activity was also characterized by the absence of LMW-ADA2 (**Fig. 8B**). Overall, we found an excellent correlation between the expression of LMW-ADA2 and extracellular (r = 0.91, p < 0.0001) or intracellular ADA2 enzyme activity (r = 0.89, p < 0.0001) (**Fig. 8C**). Two pathogenic variants lacking ADA2 enzyme activity – p.H112Q and p.T360A – showed moderate expression of LMW-ADA2. By contrast, p.P334L was characterized by absent LMW-ADA2 and a residual ADA2 enzyme activity > 25% (**Fig. 8C**). All the DADA2-associated variants previously reported to show comparatively high enzymatic activity also showed intermediate levels of LMW-ADA2. Four of those variants were reported in homozygous state in DADA2 patients (**Table 2**). Several of the variants were classified as benign by different pathogenicity scores despite having been reported in DADA2 patients (**Table 2**). Overall, the REVEL, CADD and PolyPhen scores correlated best with pathogenicity by lack of LMW-ADA2 (**Fig. 8D** and **Table S2+3**). SIFT classified the majority of variants as pathogenic despite preserved LMW-ADA2 expression and enzymatic activity while with alpha missense even many clearly pathogenic variants were categorized as benign.

**Fig. 8:**
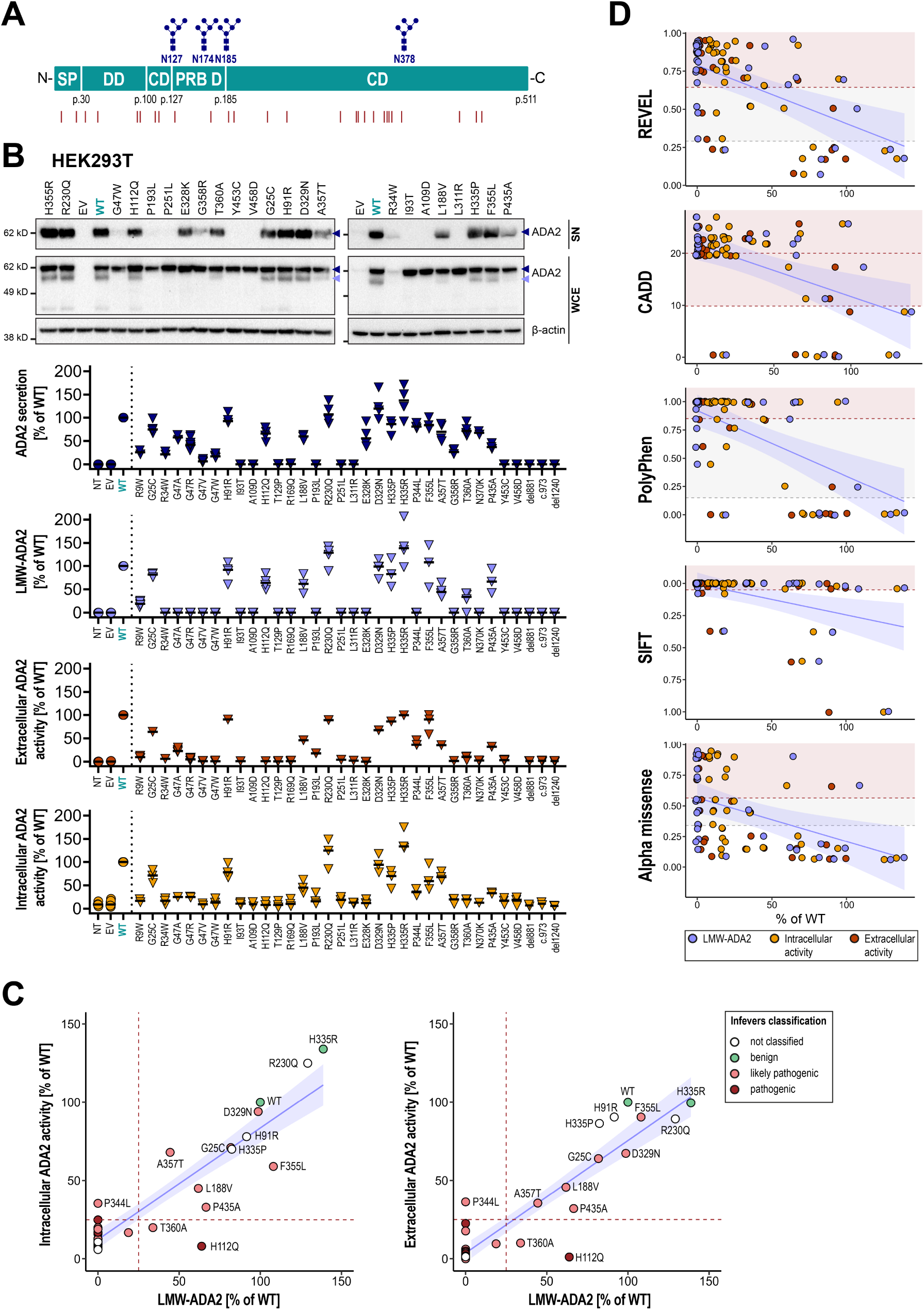
Validation of absent LMW-ADA2 as a pathogenicity marker of *ADA2* variants. **(A)** Distribution of all evaluated ADA2 variants (red lines) across the ADA2 protein domains. The N-glycosylation sites are indicated in blue. **(B)** ADA2 secretion and LMW-ADA2 expression in HEK293T cells transfected with *ADA2* variants were determined by western blot using anti-ADA2 antibody clone EPR25430-131 (#ab288296; abcam). Enzyme activity was measured in whole cell lysates and supernatants of the transfected cells. Dot plots show median. Each variant was examined in n=3-4 independent experiments. **(C)** Correlation of LMW-ADA2 and intracellular or extracellular ADA2 enzyme activity of *ADA2* variants expressed in HEK293T cells. The dots show median of n=3-4 independent experiments. The dotted lines indicate 25% expression/activity of WT ADA2. The blue line represents the fitted linear regression line with its 95% confidence interval shaded in light blue. **(D)** Correlation of LMW-ADA2 expression and ADA2 enzyme activity with different pathogenicity scores. Red shading indicates (likely) pathogenic scores while grey shading represents ambiguous scores. The blue line represents the fitted linear regression line with its 95% confidence interval shaded in light blue. The corresponding Pearson correlation coefficients are shown in Table S3. Legend: CD, catalytic domain; DD, dimerization domain; EV, empty vector; NT, non-transfected; PRB D, putative receptor binding domain; SP, signal peptide; WT, wild type.

**Table 2:** Variant characteristics. Classification of *ADA2* variants exhibiting inconclusive (> 25% of WT) levels of LMW-ADA2 expression or ADA2 enzyme activity. LMW-ADA2 expression and extracellular ADA2 enzyme activity are displayed as % of WT ADA2.

| Protein | LMW-ADA2 | ADA2<br>enzyme<br>activity | REVEL | CADD | PolyPhen | SIFT | Alpha<br>missense | Infervers | ClinVar | Homo-<br>zygous<br>patients | Hetero-<br>zygous<br>patients | Homozygous<br>individuals in<br>gnomAD |
| --- | --- | --- | --- | --- | --- | --- | --- | --- | --- | --- | --- | --- |
| p.G25C | 81.8 | 64.0 | 0.076 | 0.498 | benign (0) | tolerated<br>(0.61) | likely benign<br>(0.0748) | likely<br>pathogenic | uncertain<br>significance | 2 | 0 | - |
| p.H91R | 91.6 | 90.5 | 0.248 | 0.086 | benign<br>(0.001) | tolerated<br>(0.38) | likely benign<br>(0.0686) | not classified | uncertain<br>significance | 1 | 0 | 2 |
| p.H112Q | 63.9 | **1.1** | **0.956** | **19.6** | **probably**<br>**damaging**<br>**(0.987)** | **deleterious**<br>**(0)** | likely<br>pathogenic<br>(0.9015) | pathogenic | pathogenic/likely<br>pathogenic | 1 | 11 | - |
| p.L188V | 61.8 | 45.7 | **0.657** | **21.8** | **possibly**<br>**damaging**<br>**(0.841)** | **deleterious**<br>**(0)** | likely benign<br>(0.1571) | likely<br>pathogenic | conflicting | 0 | 2 | 2 |
| p.R230Q | 129.2 | 89.3 | 0.232 | 0.437 | benign<br>(0.007) | tolerated (1) | likely benign<br>(0.0611) | not classified | likely benign | 0 | 0 | 2 |
| p.D329N | 98.7 | 67.3 | **0.694** | **25.7** | **probably**<br>**damaging**<br>**(1)** | **deleterious**<br>**(0.02)** | likely benign<br>(0.1525) | likely<br>pathogenic | uncertain<br>significance | 1 | 0 | - |
| p.H335P | 82.4 | 86.4 | 0.175 | **11.27** | benign<br>(0.012) | **deleterious**<br>**(0)** | likely benign<br>(0.1858) | not classified | uncertain<br>significance | 0 | 1 | - |
| p.H335R | 138.8 | 99.6 | 0.176 | 8.752 | benign<br>(0.015) | **deleterious**<br>**(0.04)** | likely benign<br>(0.0727) | benign | benign | 0 | 0 | 94070 |
| p.P344L | **0** | 36.3 | 0.52 | **22.3** | **probably**<br>**damaging**<br>**(0.979)** | **deleterious**<br>**(0)** | ambiguous<br>(0.4429) | likely<br>pathogenic | NA | 0 | 3 | - |
| p.F355L | 108.1 | 90.5 | 0.502 | **17.34** | benign (0.25) | **deleterious**<br>**(0.04)** | likely<br>pathogenic<br>(0.6625) | likely<br>pathogenic | conflicting | 0 | 2 | - |
| p.A357T | 44.5 | 35.5 | **0.919** | **23.8** | **probably**<br>**damaging**<br>**(0.993)** | **deleterious**<br>**(0)** | likely benign<br>(0.2518) | likely<br>pathogenic | conflicting | 1 | 3 | - |
| p.T360A | 33.8 | **10.0** | **0.774** | **22.8** | **probably**<br>**damaging**<br>**(0.994)** | **deleterious**<br>**(0)** | ambiguous<br>(0.465) | likely<br>pathogenic | pathogenic | 10 | 15 | - |
| p.P435A | 66.6 | 32.0 | **0.706** | **22.8** | **probably**<br>**damaging**<br>**(0.997)** | **deleterious**<br>**(0)** | likely benign<br>(0.1426) | likely<br>pathogenic | likely benign | 0 | 2 | - |

In summary, we confirmed that LMW-ADA2 expression of ADA2 protein variants strongly correlates with their adenosine deaminase activity and verified absence of LMW-ADA2 as a robust indicator of pathogenicity.

## Discussion

In this study, we showed that pathogenic protein variants of ADA2 expressed in HMDM from DADA2 patients do not undergo glycan processing beyond the ER and that DADA2 HMDM therefore lack lysosomal LMW-ADA2 and secreted HMW(c)-ADA2. We reported that the LMW-form represents an intracellular glycoform of ADA2 that is generated by α-mannosidase-mediated glycan cleavage after transport to the Golgi apparatus.

The importance of N-glycosylation in the trafficking and function of ADA2 has previously been established (*12, 13*). Yet, the behavior of mutant ADA2 at the cellular level has to date only been examined in overexpression systems of cells that do not express the protein endogenously (*13, 14, 24*).

Importantly, expression of glycosyltransferases and glycosidases and therefore glycosylation patterns of individual proteins show striking variation even between different immune cell populations (*30*). It is therefore essential to understand the behavior of ADA2 in the patients’ primary cells. In our cohort of 10 DADA2 patients, we identified the intracellular predominance of unprocessed HMW-ADA2 in the absence of LMW-ADA2 as a feature that was shared by all missense variants, independently of the patients’ clinical phenotype. Differential patterns of glycosylation of the same protein being associated with distinctive protein functions have previously been described: IL-24 is a signal peptide-containing glycoprotein that is secreted via the ER. A partially glycosylated cytosolic form of IL-24 has been shown to mediate protein kinase R activation independently of its extracellular function (*31*). The predominance of the partially glycan-trimmed LMW-form of ADA2 in HC HMDM raises the question of whether, in a similar way, this form differs in function from secreted HMW(c)-ADA2. In fact, the function of ADA2 has repeatedly been subject to discussion for several reasons: (i.) Pathogenic ADA2 variants display a wide spectrum of residual adenosine deaminase function which does not clearly correlate with disease severity (*29*). (ii.) ADA2 has evolved towards lower affinity to adenosine in higher species (*11*). (iii.) ADA1 exhibits superior adenosine deaminase activity (Km of 20-50 µM vs. 2-3 mM for ADA2) and extracellular ADA1 bound to CD26 theoretically compensates for the absence of ADA2 at physiological levels of adenosine (*17*). Yet, the majority of studies proposing possible pathomechanisms underlying the immunological phenotype of DADA2 rely on the role of ADA2 as an extracellular adenosine deaminase (*15, 16*). In contrast, two groups introduced the possibility of an intracellular function of ADA2 as a lysosomal protein (*28, 32*). Greiner-Tollersrud and colleagues characterized the glycan structures of ADA2 isolated from porcine brain and identified a high mannose-6-phosphate content, indicating targeting of ADA2 to the lysosome. Indeed, they showed an overlap between the glycosylation pattern of ADA2 with several known lysosomal proteins (*28*). We showed that the glycan structures of LMW- and HMW-ADA2 expressed in HMDM were sensitive to glycan removal by Endo H, suggesting the presence of high-mannose oligosaccharides (*33*). This is in line with a potential lysosomal localization of ADA2 that we also confirmed by imaging. Moreover, we showed that multiple – also lysosomal – α-mannosidases are involved in the processing of LMW-ADA2. Combined inhibition of class I and II α-mannosidases caused an increase in the molecular weight of LMW-ADA2 yielding a glycoform almost identical to HMW-ADA2. The current lack of an antibody that sufficiently pulls down the LMW-glycoform by immunoprecipitation prevents a thorough glycan and interactome analysis of LMW-ADA2 at this point in time. We were however able to show that the predicted glycan structures derived from our inhibition experiments aligned perfectly with glycan analyses performed on porcine ADA2 (**Fig. S8A**) (*28*). Combined with the previous studies (*28, 32*), our data provide a strong hint that LMW-ADA2 represents the lysosomal glycoform of ADA2 and might be functionally distinct from secreted HMW(c)-ADA2. In addition, we show for the first time that exogenous WT ADA2 can be taken up by ADA2-deficient cells and – even more importantly – processed into the intracellular LMW-glycoform (**Fig. 7C+D**). This finding is crucial from a clinical perspective given current endeavors to develop new therapeutic approaches including the use of PEGylated ADA2. Our study provides the first hint at the capacity of externally applied ADA2 to successfully carry out also intracellular functions of ADA2.

In the HEK293T model system, transfected ADA2 was predominantly expressed as the HMW-form corresponding to secreted ADA2 in these cells, highlighting the limitations of this system to understand the behavior of endogenous ADA2. Discrepancies between the two systems are also evident from the fact that ADA2 in whole cell lysates from transfected HEK293T cells was equally sensitive to glycan removal by PNGase F or Endo H (*13*), while we observed some residual glycan structures upon Endo H treatment of ADA2 in lysates from U-937 cells and HMDM (**Fig. 5A+B**). A caveat regarding the relevance of macrophage LMW-ADA2 in the pathogenesis of DADA2 is that in primary CD14^+^ monocytes, the cells with the highest endogenous expression of ADA2, differentially glycosylated forms of ADA2 cannot be clearly distinguished (**Fig. 1A+B**). Moreover, patient monocytes – both from patients with missense variants with normal *ADA2* mRNA expression and from patients with variants that exhibit nonsense-mediated mRNA decay – show complete absence of mutant ADA2 protein expression. This implies that mutations in the *ADA2* gene might manifest differently on a cellular level depending on the expressing cell.

This notion needs to be taken into account when examining the potential induction of ER stress by pathogenic variants of ADA2. Intracellular retention of ADA2 has only been reported after overexpression in HEK293T cells (*2, 14*). While primary monocytic cells also show impaired secretion of mutant ADA2, intracellular protein levels are very low in HMDM and close to absent in CD14^+^ monocytes. In our cohort of DADA2 patients, we did not observe an increased ER stress response in CD14^+^ monocytes. This was in line with single-cell RNA-sequencing data reported by Watanabe et al. that showed downregulation of eIF2 signaling – a main pathway of the integrated stress response (*34, 35*).

Although we could not confirm that the presence of misfolded mutant ADA2 contributes to the pathophysiology of the disease by induction of an unfolded protein response, our data highlight the need for a better understanding of the folding and trafficking mechanisms of WT and mutant ADA2. We verified that the presence of pathogenic ADA2 protein variants as well as inhibition of N-glycosylation provoked the formation of intracellular protein aggregates, thereby extending previous reports (*13, 24*).

Next to Sanger sequencing of the *ADA2* gene, the measurement of serum ADA2 enzyme activity currently represents the diagnostic gold standard for DADA2. Assuming an intracellular function of the protein, it is possible that this strategy may overlook a proportion of patients. In this article, we validated the absence of LMW-ADA2 as a new characteristic of pathogenic *ADA2* variants. We showed that the pathogenic variant p.P344L exhibited complete absence of LMW-ADA2 despite a residual enzyme activity above 25% of WT ADA2. This highlights the potential benefit of additionally checking new variants for LMW-ADA2 expression in the validation process, especially when enzyme activity is inconclusive. Overall, we did however find an excellent correlation between LMW-ADA2 expression and enzyme activity, emphasizing that in most cases pathogenic variants affect both glycan processing and catalytic activity. This is in line with a similar correlation shown by Greiner-Tollersrud and colleagues in the study of the lysosomal function of ADA2 (*28*). It is important to note that both ADA2 enzyme activity and LMW-ADA2 expression are imperfect biomarkers of pathogenicity. We identified 11 DADA2-associated variants with comparatively high LMW-ADA2 expression and enzyme activity (**Table 2**). Described in patients with a DADA2-phenotype, they have all been classified as likely disease-causing. However, given our incomplete understanding of the pathophysiology of DADA2, the pathogenicity of these variants has to be considered uncertain. For p.F355L, Nihira et al. concluded from their analyses that this variant may not be pathogenic, also considering its high allele frequency. Moreover, for ambiguous variants only found in compound heterozygous state in DADA2 patients, dominant-negative effects of the other pathogenic variant also need to be taken into account (*8*). Lastly, it has to be pointed out that for the variant p.H91R that exhibits close-to-normal enzymatic activity and LMW-ADA2 expression when expressed in HEK293T cells, a homozygous patient has been described that showed strongly reduced serum ADA2 enzyme activity (*36*). In our study, we showed that – unlike HEK293T cells – CD14^+^ monocytes expressing pathogenic *ADA2* variants show absent ADA2 expression. Therefore, it is conceivable that the phenotype of certain *ADA2* variants only becomes apparent in the patients’ primary cells.

Until now, monocytes as the main producers of endogenous ADA2 have received the most attention in deciphering the pathophysiology of the disease. Considering the role of glycosylation in protein folding and the potential impact of this modification on protein function, we cannot exclude that the main ADA2 function might vary even between different immune cells. This might also account for the diverse clinical phenotype of the disease and the varying involvement of different immune cell populations. The large number of pathogenic variants and their diversity with respect to biochemical characteristics are additional factors contributing to the difficulties in delineating a single all-encompassing pathomechanism that explains the complexity of the disease.

In conclusion, we demonstrate for the first time the differential N-glycosylation of pathogenic ADA2 in patient-derived macrophages. We establish LMW-ADA2 as a hypoglycosylated non-secreted form of ADA2 in healthy controls. This LMW-ADA2 is generated by α-mannosidases after transfer to the Golgi and is absent in DADA2. Further studies will be needed to confirm the functional relevance of these results in the pathogenesis of the disease.

## Materials and Methods

### Experimental design

This exploratory study included patients treated at University Hospitals Leuven and their parents were included as heterozygous carriers. The diagnosis of DADA2 was made based on the clinical phenotype followed by a combination of serum ADA2 enzyme activity and sequencing of the *ADA2* gene according to the DADA2 management guidelines (*37*). Healthy donors were recruited at University Hospitals Leuven and KU Leuven. This study was approved by the Ethics Committee for Research of Leuven University Hospitals (project numbers S63077, S63807). The patient with α-mannosidosis was recruited at Charité University Hospital in Berlin. The inclusion of the patient from Berlin was approved by the Ethics Committee of Charité University Medicine Berlin (project number EA2/1178/22). The patients provided written informed consent prior to participation. The study was performed in accordance with the ethical standards as laid down in the 1964 Declaration of Helsinki and its later amendments. Our study examined male and female individuals, and similar findings are reported for both sexes.

The aim of this study was to determine the protein characteristics of ADA2 endogenously expressed in human immune cells carrying pathogenic *ADA2* variants. To compensate for the limited sample size in the study of patients with rare diseases, our results were validated in two separate *in vitro* models of ADA2 deficiency examining a total of 34 pathogenic *ADA2* variants. The results were confirmed using two different anti-ADA2 antibodies both validated with the help of ADA2 knockout samples and reproduced in two technically distinct blotting systems to exclude technical artefacts.

### Sanger sequencing

Genomic DNA samples were prepared from heparinized peripheral blood following the instructions of the QIAamp DNA Blood Mini kit (#51104; QIAGEN). Primers were designed with the help of Oligo Primer Analysis Software version 7 (Molecular Biology Insights). ADA2-specific gDNA amplification was performed using Platinum™ SuperFi™ PCR Master Mix (#12358010; Thermo Fisher Scientific). PCR products were purified using the QIAquick PCR purification kit (#28106; QIAGEN). Sanger sequencing was performed on an ABI 3730 XL Genetic Analyzer (Applied Biosystems) at LGC Genomics (Berlin, Germany). Sequencing data were analyzed using Chromas 2.6.5 (http://www.technelysium.com.au).

### Cell culture

Peripheral blood mononuclear cells (PBMCs) were isolated by density gradient centrifugation using Lymphoprep™ (#1114546; PROGEN) in SepMate™ isolation tubes (#85450; STEMCELL Technologies) according to the manufacturer’s instructions. Prior to monocyte isolation, the final centrifugation step was performed at 300*xg* for 10 minutes at 4 °C. CD14^+^ monocytes were isolated magnetically by positive selection using human CD14 MicroBeads (#130-050-201; Miltenyi Biotec) on LS columns (#130-042-401; Miltenyi Biotec) according to the manufacturer’s protocol. The cells were eluted into complete medium (RPMI 1640 medium (#61870044; Gibco) supplemented with 10% fetal calf serum (FCS) (#S181BH-500, Biowest) and 1% penicillin-streptomycin (#15140122; Gibco). Purity > 97% of CD14^+^ monocytes after magnetic sorting was verified by flow cytometry (anti-CD14-FITC clone MφP9, 1:40, #345784; BD Biosciences). For macrophage differentiation, 3×10^5^ CD14^+^ monocytes were seeded in a 12-well plate in 1 mL complete medium containing 20 ng/mL GM-CSF (#300-03; Peprotech) or 50 ng/mL M-CSF (#300-25; Peprotech). The cells were differentiated for ten days, with medium changes every three days. U-937, THP-1 and Jurkat cells (all purchased from ATCC) were cultured in complete RPMI medium. HEK293T cells (ATCC) were cultured in DMEM supplemented with 10% FCS and 1% penicillin-streptomycin. For transfection, the cells were seeded in medium containing no antibiotics.

Inhibition of the secretory pathway was achieved by 24-hour incubation with 1 µg/mL brefeldin A (#00-4506-51; Thermo Fisher Scientific) or 2 µg/mL monensin (GolgiStop™, #554724, BD Biosciences). For analysis of N-glycosylation and glycan trimming, the cells were incubated with tunicamycin (#T7765; Sigma-Aldrich) at 2.5 µg/mL (HEK293T cells) or 5 µg/mL (HMDM) and castanospermine (#BML-S107-0100; Enzo Life Sciences) at 100 µg/mL for 24h or 48h. α-mannosidases were inhibited by 72-96h incubation with 375 nM kifunensine (#K1140; Sigma-Aldrich) and/or 10 µM swainsonine (#S8195; Sigma-Aldrich). Monocytes were treated with 1 µg/mL tunicamycin for 24h for ER stress induction.

### Transfection

The plasmid expressing myc-DDK-tagged wild-type ADA2 (transcript variant 3, NM_001282225) was purchased from OriGene Technologies (#RC238645). Pathogenic *ADA2* variants were created by site-directed mutagenesis using the Q5® Site-Directed Mutagenesis Kit (#E0554; New England Biolabs) according to the manufacturer’s instructions. Stable competent E. coli (#C3040H, New England Biolabs) were transformed with the generated constructs. Plasmid DNA was purified with the help of the QIAprep Spin Miniprep Kit (#27104, QIAGEN). Successful mutagenesis was verified by Sanger sequencing (LGC Genomics). HEK293T cells were seeded at 5×10^4^ cells/well in 1 mL in a 24-well plate or 2.5×10^5^ cells/well in 2 mL in a 6-well plate 24h prior to transfection. The cells were transfected with 5 ng and 25 ng plasmid DNA, respectively, using Lipofectamine™ 2000 Transfection Reagent (#11668019; Invitrogen) according to the manufacturer’s instructions. The medium was changed after 24h. Cells and supernatants were collected 48h after transfection. For the reuptake experiments, HEK293T cells were transfected with 1 µg plasmid expressing untagged wild-type ADA2 or tagged wild-type and mutant as indicated. Supernatants were collected after 48 hours and transferred to U-937 cells for a 6-24-hour incubation period.

### Generation of ADA2^-/-^ cell lines by CRISPR/Cas9

Single-guide RNAs targeting ADA2 from the human CRISPR Brunello library (#73179; addgene) (*38*) were cloned into the lentiCRISPRv2 puro plasmid. lentiCRISPRv2 puro was a gift from Brett Stringer (plasmid #98290; addgene; http://n2t.net/addgene:98290; RRID: Addgene_98290). U-937 cells were transfected by electroporation using the Neon™ Transfection System (#MPK5000; Thermo Fisher Scientific) according to the manufacturer’s instructions: 2×10^5^ cells were resuspended in Resuspension Buffer R and mixed with 1 µg plasmid DNA. Electroporation was achieved with the following conditions: 1325V, 10ms, 3 pulses. After electroporation, the cells were directly transferred into 500 µL prewarmed complete RPMI 1640 medium. On day 1 after transfection, puromycin (#ant-pr-1; InvivoGen) was added at a final concentration of 1 µg/mL. After 36h, cells were seeded at 1 cell/well in 100 µL RPMI 1640 medium supplemented with 20% FCS in a round bottom 96-well plate. After 14 days, the clones were screened for knock out of ADA2 by western blot.

### Transduction

Pathogenic *ADA2* variants were created by site-directed mutagenesis as described above. Fragments containing the sequence coding for the untagged ADA2 protein were then generated using the following primers and CloneAmp™ HiFi PCR Premix (#639298; Takara Bio): ORF_ADA2_PSPXI_F_GGTGGTACTCGAGTATGTTGGTGGATGGCCCATCTGA; ORF_ADA2_PmeI_R_GGTGGTGTTTAAACCTTTGTAGCCACATCTGCTATGAACTT.

The fragments were cleaned up using the QIAquick PCR Purification Kit (#28104; QIAGEN) and cloned into the p3D lentiviral expression vector as follows: Restriction of the expression vector was performed in CutSmart buffer (#B7204; New England Biolabs) with PspX1 (#R0656; New England Biolabs) and PmeI (#R0560; New England Biolabs) at 37 °C overnight followed by 20-minute heat inactivation at 65 °C. Dephosphorylation of 5’-ends was performed using rSAP (#M0371; New England Biolabs) at 37 °C overnight followed by 20-minute heat inactivation at 65 °C. The PCR fragments were subsequently inserted into the expression vector using T4 ligase (#M0202; New England Biolabs) for 10 minutes at room temperature followed by 10-minute heat inactivation at 65 °C. After DNA cleanup (#28104; QIAGEN), stable competent E. coli (#C3040H, New England Biolabs) were transformed with the generated plasmids. Plasmid DNA was purified with the help of the QIAprep Spin Miniprep Kit (#27104, QIAGEN). Sanger sequencing (LGC Genomics) was performed to confirm that the genetic information for ADA2 was intact and that the respective mutations were present. Lentiviral particles were generated by transfecting HEK293T cells with the plasmids p3A (Gag-Pol), p3B (Rev), p3C (VSV-G) and p3D (expression vector) at a ratio of 1:1:1:3 using Lipofectamine™ 2000 Transfection Reagent (#11668019; Invitrogen). Supernatants containing lentiviral particles were harvested 48h and 72h after transfection. The viral titer was determined using the qPCR Lentivirus Titer Kit (#LV900; Applied Biological Materials). Viral supernatants were prepared at MOI=40 with 8 µg/mL polybrene and 10^6^ U-937 cells were transduced by spinoculation at 700*xg* for 2 hours at 32 °C. After 4 hours at 37 °C, the supernatant was replaced by fresh complete RPMI medium. Selection with 1 µg/mL puromycin (#ant-pr-1; InvivoGen) was started 48 hours after the transduction.

### Subcellular fractionation

Membrane and cytosolic fractions were generated from 10×10^6^ U-937 cells using the Mem-PER™ Plus Membrane Protein Extraction Kit (#89842; Thermo Fisher Scientific). Incubation times were optimized to increase protein yield to 45 minutes for the permeabilization buffer and 90 minutes for the solubilization buffer. The lysosomal fraction was produced with the help of the Lysosome Enrichment Kit for Tissues and Cultured Cells (#89839; Thermo Fisher Scientific) according to the manufacturer’s protocol.

### Glycan removal

Glycan removal was performed on whole cell lysates generated with NP-40 lysis buffer or precipitated supernatant protein reconstituted in NP-40 lysis buffer. Glycan removal was performed on 20 µg, 10 µg or 25 µg total protein using PNGase F (#P0704; New England Biolabs), Endo H (#P0702; New England Biolabs), α1-2,3,6 Mannosidase (#P0768; New England Biolabs) or PNGase F (#A39245; Gibco) according to the manufacturer’s instructions. Glycan removal with PNGase F (#P0704; New England Biolabs) was completed under denaturing conditions for 1 h at 37 °C or under non-denaturing conditions for 4 h at 37 °C as indicated.

### Immunoblotting

Adherent cells were detached with trypsin-EDTA (0.05%) (#25300062; Gibco) for 5 minutes at 37 °C and the reaction was blocked by addition of complete medium. Cells were pelleted by centrifugation at 400*xg* for 5 minutes at 4 °C. Whole cell lysates were obtained by lysing 1×10^6^ cells in 25 µL RIPA buffer (150 mM NaCl, 1% Triton X-100, 0.5% sodium deoxycholate, 0.1% SDS, pH 8.0) or NP-40 lysis buffer (150 mM NaCl, 50 mM Tris-HCl, 1% NP-40, pH 7.4) containing protease inhibitor (#78429; Thermo Fisher Scientific) for 30 minutes on ice, followed by centrifugation at 13,500*xg* for 20 minutes at 4 °C.

To collect supernatants, cells were cultured in serum-free medium for 24 h. For immunoblotting, supernatants were concentrated by adding 1200 µL ice-cold acetone to 300 µL supernatant and overnight storage at −20°C. Supernatant proteins were then precipitated by centrifugation at 15,000*xg* for 15 minutes at 4°C. The protein pellet was resuspended in 25 µL NP-40 buffer containing protease inhibitor. Bolt™ LDS sample buffer (#B0007; Thermo Fisher Scientific) mixed with Bolt™ Sample Reducing Agent (#B0009; Thermo Fisher Scientific) was added to the samples prior to gel electrophoresis. For analysis of intracellular dimer formation, the reducing agent was omitted. Samples were heated for 5 minutes at 70 °C. Supernatants from HEK293T cells transfected with ADA2 were diluted 1:10 in 2X Bolt™ LDS sample buffer (#B0007; Thermo Fisher Scientific), serum was diluted 1:200. SeeBlue™ Plus2 Pre-stained Protein Standard (#LC5925; Thermo Fisher Scientific) was used as protein molecular weight marker.

Proteins were transferred onto PVDF transfer membranes. The membranes were probed with the following primary antibodies: anti-ADA2 (clone EPR25430-131, #ab288296, 1:1000; abcam), anti-ADA2 (#HPA007888, 1:500; Sigma-Aldrich; RRID: AB_1078495), Clone OTI4C5, anti-DDK (FLAG) (clone OTI4C5, #TA50011, 1:1000; OriGene Technologies; RRID: AB_2622345), anti-BiP (#3183, 1:1000; Cell Signaling; RRID: AB_10695864), anti-LAMP-1 (clone H4A3, #sc-20011; 1:500; Santa Cruz Biotechnology; RRID: AB_626853), anti-CD98 (clone E-5, #sc-376815, 1:500; Santa Cruz Biotechnology; RRID: AB_2938854), anti-GAPDH (clone 7B, #sc-69778, 1:500; Santa Cruz Biotechnology; RRID: AB_1124759), anti-β-actin (clone: AC-15, #A5441, 1:9,000; Sigma-Aldrich; RRID: AB_476744) at 4°C overnight or at room temperature for two hours. The membranes were washed and incubated with the respective HRP-coupled secondary antibodies for one hour at room temperature: goat anti-rabbit IgG H&L (#ab205718, 1:5000; abcam; RRID: AB_2819160), goat anti-mouse IgG (H + L) (#71045, 1:5000; Sigma-Aldrich; RRID: AB_11211441) or VeriBlot Detection Reagent (#ab131366, 1:1000 in tris saline/5% skim milk; abcam).

The anti-ADA2 antibody (clone EPR25430-131, #ab288296, 1:1000; abcam) was used in the following figures of the article: 1A, 1B, 1C, 1D, 1E, 3A, 3B, 3C (left), 3D, 3E, 3F, 4D, 4E, 5B (right), 6A, 6B, 6C (top), 7C, 7D, S2C, S2D, S2E, S2F, S3A, S3D, S3F, S4, S5C, S5E, S5F, S6 and S7. The anti-ADA2 antibody (#HPA007888, 1:500; Sigma-Aldrich; RRID: AB_1078495) was applied in the remaining blots: 2B, 2C, 2D, 2E, 3C (right), 5A, 5B (left), 6C (bottom), 7B, S3B, S3C, S3E, S5B.

Protein expression was visualized by enzymatic chemiluminescence using Pierce^TM^ ECL western blotting substrate (#32106; Thermo Fisher Scientific) or SuperSignal™ West Pico PLUS Chemiluminescent Substrate (#34580; Thermo Fisher Scientific) in a ChemiDoc XRS+ Imaging System (Bio-Rad).

The experiments examining the glycan-processing of LMW-ADA2 were performed in a Wes Automated Western Blot System (ProteinSimple / bio-techne) according to the manufacturer’s instructions. Samples were loaded at a final protein concentration of 0.4 mg/mL.

### Immunoprecipitation

Whole cell lysates were prepared from HEK293T cells transfected with wild-type *ADA2* or the variant p.R169Q or HMDM from HC or DADA2 patients in NP-40 lysis buffer at a protein concentration of 1 mg/mL. Primary antibodies were added to 1000 µg lysate or 500 µg lysate, respectively, and incubated rotating at 4 °C overnight. Alternatively, 1 mL supernatant was directly incubated with the primary antibodies. After overexpression, tagged ADA2 was pulled down using 1 µg anti-DDK (FLAG) clone OTI4C5 (#TA50011, OriGene Technologies). An isotype control sample was incubated with 1 µg mouse IgG1 (#02-6100; Invitrogen) in parallel. Endogenous ADA2 was pulled down using 1 µg anti-ADA2 (clone EPR25430-131, #ab288296, abcam) with rabbit monoclonal IgG (clone EPR25A, #ab172730; abcam) used as isotype control. For immunoprecipitation, the samples were rotated with 100 µL SureBeads™ Protein G Magnetic Beads (#1614023, Bio-Rad) for 1h at room temperature. The beads were magnetized and washed four times with PBS/0.1% Tween 20. Samples for mass spectrometric analysis were washed twice with NP-40 lysis buffer and twice with PBS. For western blot, the samples were eluted from the beads in Bolt™ LDS sample buffer (#B0007; Thermo Fisher Scientific).

### Protein digest

Beads from immunoprecipitation experiments were resuspended in 20 µL denaturation buffer (6 M Urea, 2 M Thiourea, 10 mM HEPES, pH 8.0), reduced for 30 minutes at 25 °C in 12 mM dithiothreitol, followed by alkylation with 40 mM chloroacetamide for 20 minutes at 25 °C. Samples were first digested with 0.5 µg endopeptidase LysC (Wako, Osaka, Japan) for 4 h. After diluting the samples with 80 µL 50 mM ammonium bicarbonate (pH 8.5), 1 µg sequence grade trypsin (Promega) was added overnight at 25 °C. The peptide-containing supernatant was collected and acidified with formic acid (1% final concentration) to stop the digestion. Peptides were desalted and cleaned up using Stage Tip protocol (*39*). After elution with 80% acetonitrile/0.1% formic acid, samples were dried using speedvac, resolved in 3% acetonitrile/0.1% formic acid and analyzed by LC-MS/MS.

### LC-MS/MS analyses

Peptides were separated on a reversed-phase column (20 cm fritless silica microcolumns with an inner diameter of 75 µm, packed with ReproSil-Pur C18-AQ 3 µm resin (Dr. Maisch GmbH) using a 90-minute gradient with a 250 nL/min flow rate of increasing Buffer B concentration (from 2% to 60%) on a High-Performance Liquid Chromatography (HPLC) system (Thermo Fisher Scientific) and ionized using an electrospray ionization (ESI) source (Thermo Fisher Scientific) and analyzed on a Thermo Q Exactive HF-X instrument (HEK293T) or on an Q Exactive Plus instrument (HMDM). The Q Exactive HF-X instrument was run in data dependent mode selecting the top 20 most intense ions in the MS full scans, selecting ions from 350 to 2000 m/z, using 60 K resolution with a 3×10^6^ ion count target and 10 ms injection time. Tandem MS was performed at a resolution of 15 K. The MS2 ion count target was set to 1×10^5^ with a maximum injection time of 100 ms. Only precursors with charge state 2–6 were selected for MS2. The dynamic exclusion duration was set to 30 s with a 10-ppm tolerance around the selected precursor and its isotopes. The Q Exactive Plus instrument was run in data dependent mode selecting the top 10 most intense ions in the MS full scans, selecting ions from 350 to 2000 m/z, using 70 K resolution with a 3×10^6^ ion count target and 50 ms injection time. Tandem MS was performed at a resolution of 17.5 K. The MS2 ion count target was set to 5×10^4^ with a maximum injection time of 250 ms. Only precursors with charge state 2–6 were selected for MS2. The dynamic exclusion duration was set to 30 s with a 10-ppm tolerance around the selected precursor and its isotopes.

Raw data were analyzed using MaxQuant software package (v1.6.3.4) (*40*). The internal Andromeda search engine was used to search MS2 spectra against a human UniProt database (HUMAN.2020-06), including ADA2 mutant sequence, containing forward and reverse sequences. The search included variable modifications of methionine oxidation, N-terminal acetylation and fixed modification of carbamidomethyl cysteine. The FDR was set to 1% for peptide and protein identifications. Unique and razor peptides were considered for quantification. The LFQ (label-free quantitation) algorithm was activated. The resulting text file was filtered to exclude reverse database hits, potential contaminants, and proteins only identified by site.

### Microscopy

HMDM were differentiated with GM-CSF or M-CSF for 10 days in a 12-well plate as described above. Cell morphology was evaluated in the culture plate by brightfield microscopy on an EVOS M7000 Microscope Imaging System (#AMF7000; Invitrogen) using the 20X objective.

The immunofluorescence staining was performed in V-shaped 96-well plates in the dark at room temperature (RT). Cells were harvested, fixed in 4% (v/v) paraformaldehyde (Electron Microscopy Sciences) for 10 minutes, and permeabilized with PBS containing 0.1% Tween®20 (Qbiogene Inc.) for 10 minutes. To prevent nonspecific antibody binding, cells were incubated in blocking buffer (1x PBS supplemented with 5% FCS and 5 mg/mL human IgG; IgG1 66.6%, IgG2 28.5%, IgG3 2.7%, IgG4 2.2%; Grifols) for 30 minutes.

For the primary staining, an unconjugated rabbit anti-ADA2 antibody (clone EPR25430-131, #ab288296, 1:100; abcam) was applied, followed by incubation with a secondary antibody (donkey anti-rabbit AF488, 1:500, #A-21206, Invitrogen or goat anti-rabbit AF546, 1:500, #A-11035, Invitrogen). In addition, co-staining was performed using rabbit anti-GM130 AF647 (clone D6B1, 1:100, #59890, Cell Signaling Technology) in combination with either rabbit anti-LAMP1 AF488 (clone D2D11, 1:100, #58996, Cell Signaling Technology) or rabbit anti-calnexin AF555 (clone C5C9, 1:100, #23198, Cell Signaling Technology). All antibodies were diluted in 1X PBS supplemented with 5% FCS, 5 mg/mL human IgG, and 0.1% Tween®20, and incubated for 2 hours at RT. After each incubation step, cells were washed three times with PBS. Nuclear staining was performed as the final step using DAPI (1 µg/mL in PBS / 0.1% Tween®20) for 15 minutes. For imaging, stained cells were transferred onto glass slides using a Cytospin 4 centrifuge (Thermo Fisher Scientific) at 800 rpm for 3 minutes with medium acceleration. Images were acquired with a laser scanning confocal fluorescence microscope (LSM 710, Carl Zeiss).

### ADA2 enzyme activity

Adenosine deaminase 2 enzyme activity was determined in whole cell lysates from HEK293T cells overexpressing ADA2 or HMDM with and without glycan removal as well as in supernatants from transfected HEK293T cells and human serum. Deaminase activity was measured in a colorimetric assay adapted from Giusti and Galanti (*41*). Erythro-9-(2-hydroxy-3-nonyl) adenine (EHNA) (#E114; Sigma-Aldrich) was used to inhibit ADA1 activity. Triplicate measurements were performed for all samples. Enzymatic activity of pathogenic ADA2 variants overexpressed HEK293T cells was normalized to the activity of wild-type ADA2.

### qPCR

5-10×10^5^ cells were lysed in TRIzol™ Reagent (#15596018; Thermo Fisher Scientific) for 3 minutes at room temperature and homogenized before storage at −80 °C. To measure the ER stress response, HEK293T cells were seeded at 5×10^4^ cells/ml in a 24-well plate 24 h prior to transfection. TRIzol™ samples were prepared 48h after transfection. RNA was extracted using the PureLink™ RNA Mini Kit (#12183018A; Thermo Fisher Scientific) according to the manufacturer’s instruction. cDNA was generated from 20 ng RNA using the SuperScript™ VILO™ cDNA Synthesis Kit (#11754050; Thermo Fisher Scientific). Quantitative polymerase chain reaction (qPCR) analysis was performed with SsoAdvanced™ Universal SYBR® Green Supermix (#1725271; Bio-Rad Laboratories) and the following primers:

ADA2(929–1038)_F_GGCTCCGAATCAAGTTCCCC; ADA2(929–1038)_R_TAAGGCAGCTTAACGCCATCC; ADA2(1138–1257)_F_ACCAGAATCGGCCATGGA; ADA2(1138–1257)_R_CTACAGGGTGGTTCCTCAAGTCA; HSPA5_F_CAAGCAACCAAAGACGCTGGA; HSPA5_R_GCCACCCAGGTCAAACACCA; DDIT3_F_AGAGGAAGAATCAAAAATCT; DDIT3_R_AGCTCTGACTGGAATCTGGA; GAPDH_F_GTCTCCTCTGACTTCAACAGCG; GAPDH_R_ACCACCCTGTTGCTGTAGCCAA.

For all conditions, three technical replicates were measured. The experiment was run on a QuantStudio™ 3 Real-Time PCR System (Thermo Fisher Scientific) and analyzed using the QuantStudio™ Design & Analysis Software v1.5.2. The relative abundance of *ADA2* was normalized to the expression level of *GAPDH*. and different conditions were compared using the 2^-ΔΔCt^ method (*42*).

### Statistical analysis

Statistical analysis was performed in R using the packages stats and Hmisc. Correlation was measured using Pearson’s correlation coefficient.

Statistical analysis of the mass spectrometry data was performed using Perseus software (v1.6.2.1) (*43*). Log2 transformed LFQ intensity values were filtered for minimum of 3 valid values in at least one experimental group and missing values were imputed with random low intensity values taken from a normal distribution. Differences in protein abundance between groups were calculated using two-sample Student’s t-test using an FDR-based significance cut-off of 5% (*HEK293T*) or 10% (*HMDM*) in order to define enriched proteins.

## Abbreviations

ADA2: Adenosine deaminase 2
BFA: Brefeldin A
DADA2: Deficiency of adenosine deaminase 2
Endo: H Endoglycosidase H
ER: Endoplasmic reticulum
GM-CSF: Granulocyte–macrophage colony-stimulating factor
HC: Healthy control
HMDM: Human monocyte-derived macrophages
HMW: High-molecular-weight
LMW: Low-molecular-weight
M-CSF: Macrophage colony-stimulating factor
PNGase F: Peptide-N-Glycosidase F
TNFi: TNF inhibitors
WT: Wild type

## Acknowledgments

We thank Prof. Jaak Jaeken, Prof. Daisy Rymen and Dr. Eva Morava-Kozicz for kindly sharing their expertise and insight on protein glycosylation. We thank Lotte Bral for her assistance in the production of the knock-out cell lines. Above all, we would like to express our gratitude to our patients and their parents for their willingness to participate in our study and to their treating physicians and nurses for their efforts in the collection of research samples. The figures Fig. 2A, Fig. 3G, Fig. 5C, Fig. 6E, Fig. 7E, Fig. 8A and Fig. S8A were created with BioRender.com.

## Funding

This project has received funding from the following sources:

European Research Council (ERC) under the European Union’s Horizon 2020 research and innovation program GA No. 948959 (MORE2ADA2) (LE, MW, BP, MD, IM)

KU Leuven C1 Grant C16/18/007 (IM)

Research Foundation-Flanders (FWO) grant G0B5120N (IM) Jeffrey Modell Foundation (IM)

ERN-RITA (IM)

Research Foundation – Flanders (FWO) grant 11E0123N (LE)

Charité – Universitätsmedizin Berlin and the Berlin Institute of Health BIH Charité Junior Clinician Scientist Fellowship (LE)

Research Foundation – Flanders (FWO) grant 11F4421N (SD)

Research Foundation – Flanders (FWO) grant 1805523N (RS)

Research Foundation – Flanders (FWO) grant G0A3320N (PA)

KU Leuven C14/21/095 InterAction consortium (PA)

EOS MetaNiche consortium N° 40007532 (PA)

iBOF/21/053 ATLANTIS network (PA)

EOS DECODE consortium N° 30837538 (PA)

## Author contributions

Conceptualization and design: LE, LM, IM

Material preparation and data collection: LE, AH, MW, BP, SD, MD, EFR, AD, JN, MK, LM

Data analysis: LE, AH, MB, MK, LM

Generation of ADA2 knock-out cell lines: LE, MJ, DD

Proteomics analyses: MK, PM, MFM, TM

Design of trafficking experiments: LE, LM, IM, FE

Design of ER stress experiments: LE; LM, IM, PA

Collection of clinical data and patient materials: LE, SD, GB, LDS, RS, SV, DC, IM

Supervision: IM

Writing – original draft: LE

Writing – review & editing: all authors

## Data and materials availability

All data needed to evaluate the conclusions of our study are provided in the manuscript or the Supplementary Materials. The mass spectrometry data have been deposited to the ProteomeXchange Consortium via the PRIDE partner repository with the dataset identifier PXD046128. The authors provide a supplementary file containing the original blots of all the presented data.

## References

1. Q. Zhou, D. Yang, A. K. Ombrello, A. V. Zavialov, C. Toro, A. V. Zavialov, D. L. Stone, J. J. Chae, S. D. Rosenzweig, K. Bishop, K. S. Barron, H. S. Kuehn, P. Hoffmann, A. Negro, W. L. Tsai, E. W. Cowen, W. Pei, J. D. Milner, C. Silvin, T. Heller, D. T. Chin, N. J. Patronas, J. S. Barber, C.-C. R. Lee, G. M. Wood, A. Ling, S. J. Kelly, D. E. Kleiner, J. C. Mullikin, N. J. Ganson, H. H. Kong, S. Hambleton, F. Candotti, M. M. Quezado, K. R. Calvo, H. Alao, B. K. Barham, A. Jones, J. F. Meschia, B. B. Worrall, S. E. Kasner, S. S. Rich, R. Goldbach-Mansky, M. Abinun, E. Chalom, A. C. Gotte, M. Punaro, V. Pascual, J. W. Verbsky, T. R. Torgerson, N. G. Singer, T. R. Gershon, S. Ozen, O. Karadag, T. A. Fleisher, E. F. Remmers, S. M. Burgess, S. L. Moir, M. Gadina, R. Sood, M. S. Hershfield, M. Boehm, D. L. Kastner, I. Aksentijevich, Early-onset stroke and vasculopathy associated with mutations in ADA2. N. Engl. J. Med. 370, 911–920 (2014).

2. P. Navon Elkan, S. B. Pierce, R. Segel, T. Walsh, J. Barash, S. Padeh, A. Zlotogorski, Y. Berkun, J. J. Press, M. Mukamel, I. Voth, P. J. Hashkes, L. Harel, V. Hoffer, E. Ling, F. Yalcinkaya, O. Kasapcopur, M. K. Lee, R. E. Klevit, P. Renbaum, A. Weinberg-Shukron, E. F. Sener, B. Schormair, S. Zeligson, D. Marek-Yagel, T. M. Strom, M. Shohat, A. Singer, A. Rubinow, E. Pras, J. Winkelmann, M. Tekin, Y. Anikster, M.-C. King, E. Levy-Lahad, Mutant adenosine deaminase 2 in a polyarteritis nodosa vasculopathy. N. Engl. J. Med. 370, 921–931 (2014).

3. I. Meyts, I. Aksentijevich, Deficiency of Adenosine Deaminase 2 (DADA2): Updates on the Phenotype, Genetics, Pathogenesis, and Treatment. J. Clin. Immunol. 38, 569–578 (2018).

4. K. J. Karczewski, L. C. Francioli, G. Tiao, B. B. Cummings, J. Alföldi, Q. Wang, R. L. Collins, K. M. Laricchia, A. Ganna, D. P. Birnbaum, L. D. Gauthier, H. Brand, M. Solomonson, N. A. Watts, D. Rhodes, M. Singer-Berk, E. M. England, E. G. Seaby, J. A. Kosmicki, R. K. Walters, K. Tashman, Y. Farjoun, E. Banks, T. Poterba, A. Wang, C. Seed, N. Whiffin, J. X. Chong, K. E. Samocha, E. Pierce-Hoffman, Z. Zappala, A. H. O’Donnell-Luria, E. V. Minikel, B. Weisburd, M. Lek, J. S. Ware, C. Vittal, I. M. Armean, L. Bergelson, K. Cibulskis, K. M. Connolly, M. Covarrubias, S. Donnelly, S. Ferriera, S. Gabriel, J. Gentry, N. Gupta, T. Jeandet, D. Kaplan, C. Llanwarne, R. Munshi, S. Novod, N. Petrillo, D. Roazen, V. Ruano-Rubio, A. Saltzman, M. Schleicher, J. Soto, K. Tibbetts, C. Tolonen, G. Wade, M. E. Talkowski, B. M. Neale, M. J. Daly, D. G. MacArthur, The mutational constraint spectrum quantified from variation in 141,456 humans. Nature 581, 434–443 (2020).

5. Infevers: an online database for autoinflammatory mutations. Copyright. Available at https://infevers.umai-montpellier.fr/ Accessed 14/08/2023. (available at https://infevers.umai-montpellier.fr/web/instructions_for_use.php).

6. P. Y. Lee, E. S. Kellner, Y. Huang, E. Furutani, Z. Huang, W. Bainter, M. F. Alosaimi, K. Stafstrom, C. D. Platt, T. Stauber, S. Raz, I. Tirosh, A. Weiss, M. B. Jordan, C. Krupski, D. Eleftheriou, P. Brogan, A. Sobh, Z. Baz, G. Lefranc, C. Irani, S. S. Kilic, R. El-Owaidy, M. R. Lokeshwar, P. Pimpale, R. Khubchandani, E. P. Chambers, J. Chou, R. S. Geha, P. A. Nigrovic, Q. Zhou, Genotype and functional correlates of disease phenotype in deficiency of adenosine deaminase 2 (DADA2). J. Allergy Clin. Immunol. 145, 1664–1672.e10 (2020).

7. M. Dzhus, L. Ehlers, M. Wouters, K. Jansen, R. Schrijvers, L. De Somer, S. Vanderschueren, M. Baggio, L. Moens, B. Verhaaren, R. Lories, G. Bucciol, I. Meyts, A Narrative Review of the Neurological Manifestations of Human Adenosine Deaminase 2 Deficiency. J. Clin. Immunol. (2023), doi:10.1007/s10875-023-01555-y.

8. M. Wouters, L. Ehlers, W. Van Eynde, M. E. Kars, S. Delafontaine, V. Kienapfel, M. Dzhus, R. Schrijvers, P. De Haes, S. Struyf, G. Bucciol, Y. Itan, A. Bolze, A. Voet, A. Hombrouck, L. Moens, B. Ogunjimi, I. Meyts, Dominant negative ADA2 mutations cause ADA2 deficiency in heterozygous carriers. J. Exp. Med. 222, e20250499 (2025).

9. N. T. Deuitch, D. Yang, P. Y. Lee, X. Yu, N. S. Moura, O. Schnappauf, A. K. Ombrello, D. Stone, H. S. Kuehn, S. D. Rosenzweig, P. Hoffmann, C. Cudrici, D. M. Levy, E. Kessler, J. B. Soep, A. D. Hay, A. Dalrymple, Y. Zhang, L. Sun, Q. Zhang, X. Tang, Y. Wu, K. Rao, H. Li, H. Luo, Y. Zhang, J. M. Burnham, M. Boehm, K. Barron, D. L. Kastner, I. Aksentijevich, Q. Zhou, TNF inhibition in vasculitis management in adenosine deaminase 2 deficiency (DADA2). J. Allergy Clin. Immunol. 149, 1812–1816.e6 (2022).

10. H. Hashem, A. R. Kumar, I. Müller, F. Babor, R. Bredius, J. Dalal, A. P. Hsu, S. M. Holland, D. D. Hickstein, S. Jolles, R. Krance, G. Sasa, M. Taskinen, M. Koskenvuo, J. Saarela, J. van Montfrans, K. Wilson, B. Bosch, L. Moens, M. Hershfield, I. Meyts, Hematopoietic stem cell transplantation rescues the hematological, immunological, and vascular phenotype in DADA2. Blood 130, 2682–2688 (2017).

11. A. V. Zavialov, X. Yu, D. Spillmann, G. Lauvau, A. V. Zavialov, Structural basis for the growth factor activity of human adenosine deaminase ADA2. J. Biol. Chem. 285, 12367–12377 (2010).

12. P. Y. Lee, Y. Huang, Q. Zhou, O. Schnappauf, M. S. Hershfield, Y. Li, N. J. Ganson, N. Sampaio Moura, O. M. Delmonte, S. S. Stone, M. J. Rivkin, S.-Y. Pai, T. Lyons, R. P. Sundel, V. W. Hsu, L. D. Notarangelo, I. Aksentijevich, P. A. Nigrovic, Disrupted N-linked glycosylation as a disease mechanism in deficiency of ADA2. J. Allergy Clin. Immunol. 142, 1363–1365.e8 (2018).

13. M. Ito, Y. Maejima, K. Nishimura, Y. Nakae, A. Ono, S. Iwaki-Egawa, A role for N-glycosylation in active adenosine deaminase 2 production. Biochim. Biophys. Acta BBA - Gen. Subj. 1866, 130237 (2022).

14. L. Chen, A. Mamutova, A. Kozlova, E. Latysheva, F. Evgeny, T. Latysheva, K. Savostyanov, A. Pushkov, I. Zhanin, E. Raykina, M. Kurnikova, I. Mersiyanova, C. D. Platt, H. Jee, K. Brodeur, Y. Du, M. Liu, A. Weiss, G. S. Schulert, J. Rodriguez-Smith, M. S. Hershfield, I. Aksentijevich, Q. Zhou, P. A. Nigrovic, A. Shcherbina, E. Alexeeva, P. Y. Lee, Comparison of disease phenotypes and mechanistic insight on causal variants in patients with DADA2. J. Allergy Clin. Immunol. (2023), doi:10.1016/j.jaci.2023.04.014.

15. C. Carmona-Rivera, S. S. Khaznadar, K. W. Shwin, J. A. Irizarry-Caro, L. J. O’Neil, Y. Liu, K. A. Jacobson, A. K. Ombrello, D. L. Stone, W. L. Tsai, D. L. Kastner, I. Aksentijevich, M. J. Kaplan, P. C. Grayson, Deficiency of adenosine deaminase 2 triggers adenosine-mediated NETosis and TNF production in patients with DADA2. Blood 134, 395–406 (2019).

16. R. Dhanwani, M. Takahashi, I. T. Mathews, C. Lenzi, A. Romanov, J. D. Watrous, B. Pieters, C. C. Hedrick, C. A. Benedict, J. Linden, R. Nilsson, M. Jain, S. Sharma, Cellular sensing of extracellular purine nucleosides triggers an innate IFN-β response. Sci. Adv. 6, eaba3688 (2020).

17. T. K. Tarrant, S. J. Kelly, M. S. Hershfield, Elucidating the pathogenesis of adenosine deaminase 2 deficiency: current status and unmet needs. Expert Opin. Orphan Drugs 9, 257–264 (2021).

18. O. K. Greiner-Tollersrud, V. Boehler, E. Bartok, M. Krausz, A. Polyzou, J. Schepp, M. Seidl, J. O. Olsen, C. R. Smulski, S. Raieli, K. Hübscher, E. Trompouki, R. Link, H. Ebersbach, H. Srinivas, M. Marchant, D. Staab, D. Guerini, S. Baasch, P. Henneke, G. Kochs, G. Hartmann, R. Geiger, B. Grimbacher, M. Warncke, M. Proietti, ADA2 is a lysosomal DNase regulating the type-I interferon response (2020; https://www.biorxiv.org/content/10.1101/2020.06.21.162990v2), p. 2020.06.21.162990.

19. L. Ehlers, M. Wouters, B. Pillay, S. Delafontaine, M. Dzhus, M. Baggio, T. Niehues, G. Dückers, L. Sevenants, K. Casteels, L. D. Somer, R. Schrijvers, S. Vanderschueren, M. Jacquemyn, D. Daelemans, A. Hombrouck, E. P. Chambers, T. Tousseyn, G. Bucciol, P. Agostinis, L. Moens, I. Meyts, Inhibition of lysosomal degradation increases expression of mutant ADA2 in DADA2 monocytes. J. Allergy Clin. Immunol. 0 (2025), doi:10.1016/j.jaci.2025.06.009.

20. A. Varki, R. D. Cummings, J. D. Esko, P. Stanley, G. W. Hart, M. Aebi, D. Mohnen, T. Kinoshita, N. H. Packer, J. H. Prestegard, R. L. Schnaar, P. H. Seeberger, Eds., Essentials of Glycobiology (Cold Spring Harbor Laboratory Press, Cold Spring Harbor (NY), ed. 4th, 2022; http://www.ncbi.nlm.nih.gov/books/NBK579918/).

21. T. Fujiwara, K. Oda, S. Yokota, A. Takatsuki, Y. Ikehara, Brefeldin A causes disassembly of the Golgi complex and accumulation of secretory proteins in the endoplasmic reticulum. J. Biol. Chem. 263, 18545–18552 (1988).

22. H. H. Mollenhauer, D. James Morré, L. D. Rowe, Alteration of intracellular traffic by monensin; mechanism, specificity and relationship to toxicity. Biochim. Biophys. Acta BBA - Rev. Biomembr. 1031, 225–246 (1990).

23. P. Kirchner, M. Bourdenx, J. Madrigal-Matute, S. Tiano, A. Diaz, B. A. Bartholdy, B. Will, A. M. Cuervo, Proteome-wide analysis of chaperone-mediated autophagy targeting motifs. PLOS Biol. 17, e3000301 (2019).

24. S. M. Bowers, M. Sundqvist, P. Dancey, D. A. Cabral, K. L. Brown, Pathogenic variant c.1052T>A (p.Leu351Gln) in adenosine deaminase 2 impairs secretion and elevates type I IFN responsive gene expression. Front. Immunol. 13 (2022) (available at https://www.frontiersin.org/articles/10.3389/fimmu.2022.995191).

25. K. Legler, R. Rosprim, T. Karius, K. Eylmann, M. Rossberg, R. M. Wirtz, V. Müller, I. Witzel, B. Schmalfeldt, K. Milde-Langosch, L. Oliveira-Ferrer, Reduced mannosidase MAN1A1 expression leads to aberrant N-glycosylation and impaired survival in breast cancer. Br. J. Cancer 118, 847–856 (2018).

26. D. R. Rose, Structure, mechanism and inhibition of Golgi α-mannosidase II. Curr. Opin. Struct. Biol. 22, 558–562 (2012).

27. D. Malm, Ø. Nilssen, Alpha-mannosidosis. Orphanet J. Rare Dis. 3, 21 (2008).

28. O. K. Greiner-Tollersrud, M. Krausz, V. Boehler, A. Polyzou, M. Seidl, A. Spahiu, Z. Abdullah, K. Andryka-Cegielski, F. I. Dominick, K. Huebscher, A. Goschin, C. R. Smulski, E. Trompouki, R. Link, H. Ebersbach, H. Srinivas, M. Marchant, G. Sogkas, D. Staab, C. Vågbø, D. Guerini, S. Baasch, E. Latz, G. Hartmann, P. Henneke, R. Geiger, X. P. Peng, B. Grimbacher, E. Bartok, I. Alseth, M. Warncke, M. Proietti, ADA2 is a lysosomal deoxyadenosine deaminase acting on DNA involved in regulating TLR9-mediated immune sensing of DNA. Cell Rep. 43 (2024), doi:10.1016/j.celrep.2024.114899.

29. H. Jee, Z. Huang, S. Baxter, Y. Huang, M. L. Taylor, L. A. Henderson, S. Rosenzweig, A. Sharma, E. P. Chambers, M. S. Hershfield, Q. Zhou, F. Dedeoglu, I. Aksentijevich, P. A. Nigrovic, A. O’Donnell-Luria, P. Y. Lee, Comprehensive analysis of ADA2 genetic variants and estimation of carrier frequency driven by a function-based approach. J. Allergy Clin. Immunol. (2021), doi:10.1016/j.jaci.2021.04.034.

30. M. A. Daniels, K. A. Hogquist, S. C. Jameson, Sweet “n” sour: the impact of differential glycosylation on T cell responses. Nat. Immunol. 3, 903–910 (2002).

31. S. Davidson, C.-H. Yu, A. Steiner, F. Ebstein, P. J. Baker, V. Jarur-Chamy, K. Hrovat Schaale, P. Laohamonthonkul, K. Kong, D. J. Calleja, C. R. Harapas, K. R. Balka, J. Mitchell, J. T. Jackson, N. D. Geoghegan, F. Moghaddas, K. L. Rogers, K. D. Mayer-Barber, A. A. De Jesus, D. De Nardo, B. T. Kile, A. J. Sadler, M. C. Poli, E. Krüger, R. Goldbach Mansky, S. L. Masters, Protein kinase R is an innate immune sensor of proteotoxic stress via accumulation of cytoplasmic IL-24. Sci. Immunol. 7, eabi6763 (2022).

32. L. Dong, W. Luo, S. Maksym, S. C. Robson, A. V. Zavialov, Adenosine deaminase 2 regulates the activation of the toll-like receptor 9 in response to nucleic acids. Front. Med. (2024), doi:10.1007/s11684-024-1067-5.

33. M.-S. Kim, D. Leahy, in Methods in Enzymology, Laboratory Methods in Enzymology: Cell, Lipid and Carbohydrate. J. Lorsch, Ed. (Academic Press, 2013), vol. 533, pp. 259–263.

34. N. Watanabe, S. Gao, Z. Wu, S. Batchu, S. Kajigaya, C. Diamond, L. Alemu, D. Q. Raffo, P. Hoffmann, D. Stone, A. K. Ombrello, N. S. Young, Analysis of deficiency of adenosine deaminase 2 pathogenesis based on single-cell RNA sequencing of monocytes. J. Leukoc. Biol. 110, 409–424 (2021).

35. K. Pakos-Zebrucka, I. Koryga, K. Mnich, M. Ljujic, A. Samali, A. M. Gorman, The integrated stress response. EMBO Rep. 17, 1374–1395 (2016).

36. S. Cooray, E. Omyinmi, Y. Hong, C. Papadopoulou, L. Harper, E. Al-Abadi, R. Goel, S. Dubey, M. Wood, S. Jolles, S. Berg, M. Ekelund, K. Armon, D. Eleftheriou, P. A. Brogan, Anti-tumour necrosis factor treatment for the prevention of ischaemic events in patients with deficiency of adenosine deaminase 2 (DADA2). Rheumatol. Oxf. Engl. 60, 4373–4378 (2021).

37 P. Y. Lee, B. A. Davidson, R. S. Abraham, B. Alter, J. I. Arostegui, K. Bell, A. Belot, J. R. E. Bergerson, T. J. Bernard, P. A. Brogan, Y. Berkun, N. T. Deuitch, D. Dimitrova, S. A. Georgin-Lavialle, M. Gattorno, B. Grimbacher, H. Hashem, M. S. Hershfield, R. N. Ichord, K. Izawa, J. A. Kanakry, R. P. Khubchandani, F. C. C. Klouwer, E. A. Luton, A. W. Man, I. Meyts, J. M. Van Montfrans, S. Ozen, J. Saarela, G. C. Santo, A. Sharma, A. Soldatos, R. Sparks, T. R. Torgerson, I. L. Uriarte, T. A. B. Youngstein, Q. Zhou, I. Aksentijevich, D. L. Kastner, E. P. Chambers, A. K. Ombrello, DADA2 Foundation, Evaluation and Management of Deficiency of Adenosine Deaminase 2: An International Consensus Statement. JAMA Netw. Open 6, e2315894 (2023).

37. J. G. Doench, N. Fusi, M. Sullender, M. Hegde, E. W. Vaimberg, K. F. Donovan, I. Smith, Z. Tothova, C. Wilen, R. Orchard, H. W. Virgin, J. Listgarten, D. E. Root, Optimized sgRNA design to maximize activity and minimize off-target effects of CRISPR-Cas9. Nat. Biotechnol. 34, 184–191 (2016).

38. J. Rappsilber, Y. Ishihama, M. Mann, Stop and go extraction tips for matrix-assisted laser desorption/ionization, nanoelectrospray, and LC/MS sample pretreatment in proteomics. Anal. Chem. 75, 663–670 (2003).

39. S. Tyanova, T. Temu, J. Cox, The MaxQuant computational platform for mass spectrometry-based shotgun proteomics. Nat Protoc 11, 2301–2319 (2016).

40. J. Illingworth, Methods of enzymatic analysis: Third edition: Editor-in-Chief: Hans Ulrich Bergmeyer. Verlag Chemie, 1983 (vols I–III), 1984 (vols IV & V) DM258 each volume or DM2240 vols I–X inclusive. Biochem. Educ. 13, 38–38 (1985).

41. K. J. Livak, T. D. Schmittgen, Analysis of relative gene expression data using real-time quantitative PCR and the 2(-Delta Delta C(T)) Method. Methods San Diego Calif 25, 402–408 (2001).

42. S. Tyanova, T. Temu, P. Sinitcyn, A. Carlson, M. Y. Hein, T. Geiger, M. Mann, J. Cox, The Perseus computational platform for comprehensive analysis of (prote)omics data. Nat. Methods 13, 731–740 (2016).

43. A. Sharma, G. Naidu, V. Sharma, S. Jha, A. Dhooria, V. Dhir, P. Bhatia, V. Sharma, S. Bhattad, K. G. Chengappa, V. Gupta, D. P. Misra, P. P. Chavan, S. Malaviya, R. Dudam, B. Sharma, S. Kumar, R. Bhojwani, P. Gupta, V. Agarwal, K. Sharma, M. Singhal, M. Rathi, R. Nada, R. W. Minz, V. Chaturvedi, A. Aggarwal, R. Handa, A. Grossi, M. Gattorno, Z. Huang, J. Wang, R. Jois, V. S. Negi, R. Khubchandani, S. Jain, J. I. Arostegui, E. P. Chambers, M. S. Hershfield, I. Aksentijevich, Q. Zhou, P. Y. Lee, Deficiency of Adenosine Deaminase 2 in Adults and Children: Experience From India. Arthritis Rheumatol. Hoboken NJ 73, 276–285 (2021).

44. R. Caorsi, F. Penco, A. Grossi, A. Insalaco, A. Omenetti, M. Alessio, G. Conti, F. Marchetti, P. Picco, A. Tommasini, S. Martino, C. Malattia, R. Gallizzi, R. A. Podda, A. Salis, F. Falcini, F. Schena, F. Garbarino, A. Morreale, M. Pardeo, C. Ventrici, C. Passarelli, Q. Zhou, M. Severino, C. Gandolfo, G. Damonte, A. Martini, A. Ravelli, I. Aksentijevich, I. Ceccherini, M. Gattorno, ADA2 deficiency (DADA2) as an unrecognised cause of early onset polyarteritis nodosa and stroke: a multicentre national study. Ann. Rheum. Dis. 76, 1648–1656 (2017).

45. J. Schepp, M. Proietti, N. Frede, M. Buchta, K. Hübscher, J. Rojas Restrepo, S. Goldacker, K. Warnatz, J. Pachlopnik Schmid, A. Duppenthaler, V. Lougaris, I. Uriarte, S. Kelly, M. Hershfield, B. Grimbacher, Screening of 181 Patients With Antibody Deficiency for Deficiency of Adenosine Deaminase 2 Sheds New Light on the Disease in Adulthood. Arthritis Rheumatol. Hoboken NJ 69, 1689–1700 (2017).

46. S. Özen, E. D. Batu, E. Z. Taşkıran, H. A. Özkara, Ş. Ünal, N. Güleray, A. Erden, Ö. Karadağ, F. Gümrük, M. Çetin, H. E. Sönmez, Y. Bilginer, D. Ç. Ayvaz, I. Tezcan, A Monogenic Disease with a Variety of Phenotypes: Deficiency of Adenosine Deaminase 2. J. Rheumatol. 47, 117–125 (2020).

47. A. Betrains, F. Staels, L. Moens, S. Delafontaine, M. S. Hershfield, D. Blockmans, A. Liston, S. Humblet-Baron, I. Meyts, R. Schrijvers, S. Vanderschueren, Diagnosis of deficiency of adenosine deaminase type 2 in adulthood. Scand. J. Rheumatol., 1–4 (2021).

48. O. Akgun-Dogan, P. O. Simsek-Kiper, E. Taskiran, C. Lissewski, J. Brinkmann, D. Schanze, R. Göçmen, D. Cagdas, Y. Bilginer, G. E. Utine, M. Zenker, S. Ozen, İ. Tezcan, M. Alikasifoglu, K. Boduroğlu, ADA2 deficiency in a patient with Noonan syndrome-like disorder with loose anagen hair: The co-occurrence of two rare syndromes. Am. J. Med. Genet. A. 179, 2474–2480 (2019).

49. M. Rama, C. Duflos, I. Melki, D. Bessis, A. Bonhomme, H. Martin, D. Doummar, S. Valence, D. Rodriguez, E. Carme, D. Genevieve, K. Heimdal, A. Insalaco, N. Franck, V. Queyrel-Moranne, N. Tieulie, J. London, F. Uettwiller, S. Georgin-Lavialle, A. Belot, I. Koné-Paut, V. Hentgen, G. Boursier, I. Touitou, G. Sarrabay, A decision tree for the genetic diagnosis of deficiency of adenosine deaminase 2 (DADA2): a French reference centres experience. Eur. J. Hum. Genet. EJHG 26, 960–971 (2018).

50. A. K. Ombrello, J. Qin, P. M. Hoffmann, P. Kumar, D. Stone, A. Jones, T. Romeo, B. Barham, G. Pinto-Patarroyo, C. Toro, A. Soldatos, Q. Zhou, N. Deuitch, I. Aksentijevich, S. L. Sheldon, S. Kelly, A. Man, K. Barron, M. Hershfield, W. A. Flegel, D. L. Kastner, Treatment Strategies for Deficiency of Adenosine Deaminase 2. N. Engl. J. Med. 380, 1582–1584 (2019).

51. K. M. Gibson, K. A. Morishita, P. Dancey, P. Moorehead, B. Drögemöller, X. Han, J. Graham, R. E. W. Hancock, D. Foell, S. Benseler, R. Luqmani, R. S. M. Yeung, S. Shenoi, M. Bohm, A. M. Rosenberg, C. J. Ross, D. A. Cabral, K. L. Brown, Identification of Novel Adenosine Deaminase 2 Gene Variants and Varied Clinical Phenotype in Pediatric Vasculitis. Arthritis Rheumatol. Hoboken NJ 71, 1747–1755 (2019).

