## Supplemental material for "Human ADA2 deficiency is characterized by the absence of an intracellular hypoglycosylated form of adenosine deaminase 2"

Lisa Ehlers *et al.*

**This PDF file includes:**

Figures S1 to S8

Table S3

Legend for Tables S1 to S2

References (44 to 52)

**Other Supplementary Materials for this manuscript include the following:**

Tables S1 to S2

Uncropped western blot images

### Figure S1

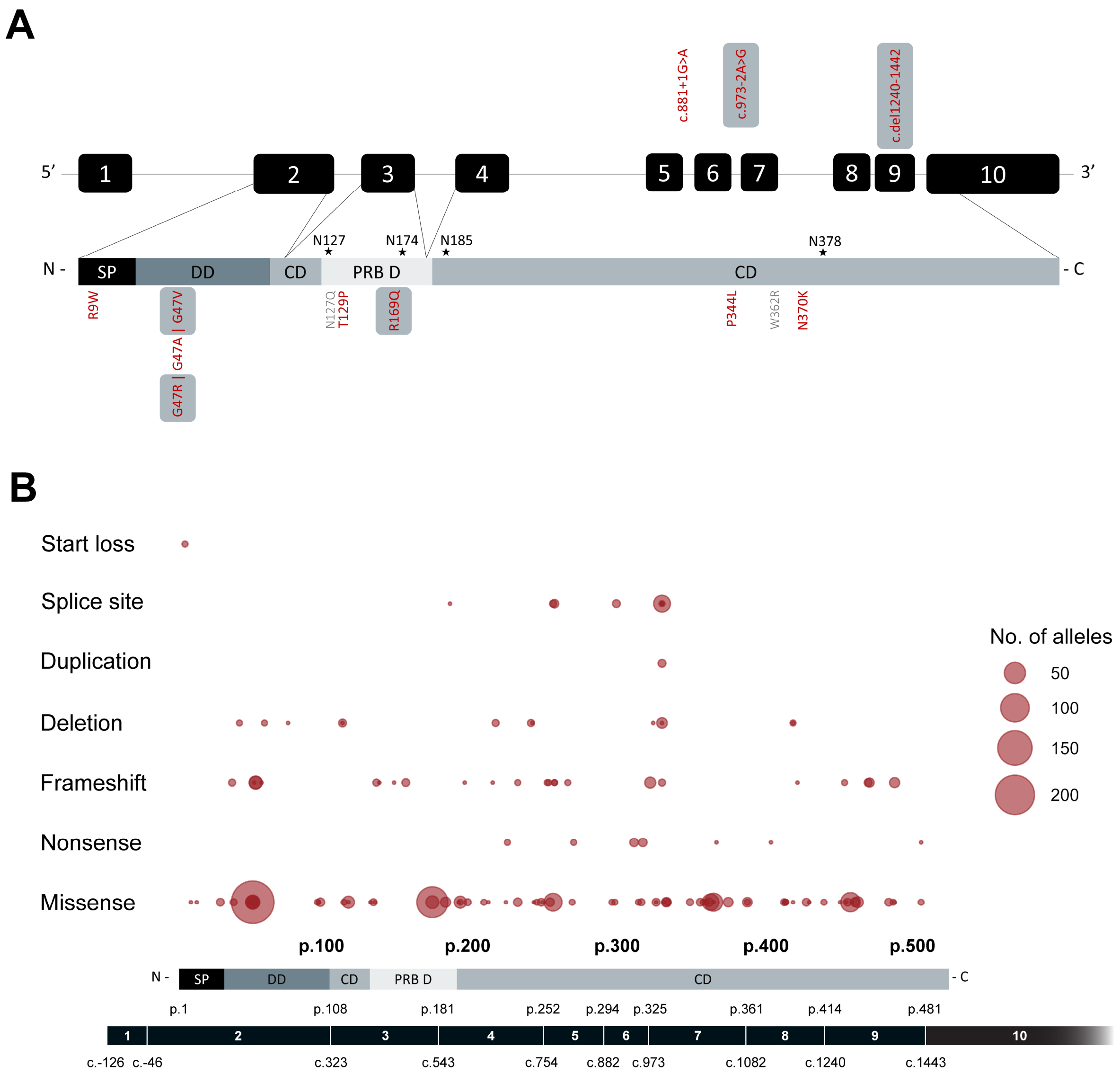


**Figure S1: Pathogenic variants in *ADA2* underlying ADA2 deficiency. (A)** Overview of the location of pathogenic variants of *ADA2* included in this study across the *ADA2* gene and encoded protein domains. Shaded variants are present in our cohort of DADA2 patients. Stars indicate N-glycosylation sites. **(B)** Overview of frequency, location and mutation type of all published pathogenic *ADA2* variants as reported by Dzhus and colleagues (*7*). Legend: CD, catalytic domain; DD, dimerization domain; EV, empty vector; PRB D, putative receptor binding domain.

### Figure S2

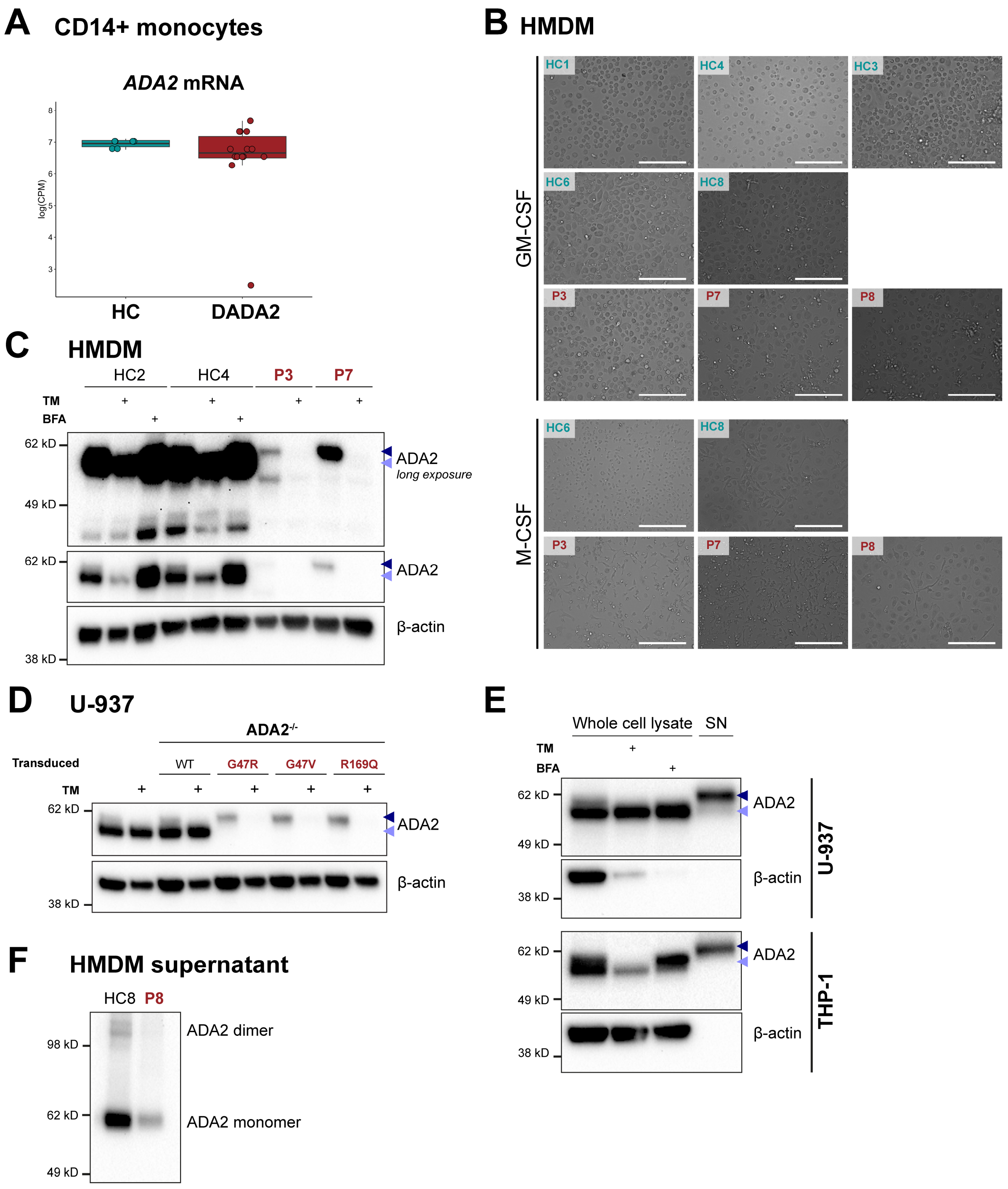


**Figure S2: Expression of ADA2 in primary immune cells and cell lines. (A)** *ADA2* mRNA levels in sorted CD14$^{+}$ monocytes. Single-cell RNA sequencing was performed on magnetically sorted CD14$^{+}$ monocytes from healthy controls (HC) and DADA2 patients (P). The graph was generated based on the data set published by Watanabe and colleagues (*34*). Boxplots show median and interquartile range, whiskers extend from the hinge to the highest and lowest value within the 1.5 * interquartile range of the hinge. **(B)** Morphology of DADA2 human monocyte-derived macrophages. $CD14^+$ monocytes from healthy controls (HC) and DADA2 patients (P) were differentiated over ten days with 20 ng/mL GM-CSF or 50 ng/mL M-CSF and imaged by brightfield microscopy on day 10. The scale bar represents 200 μM. **(C)** ADA2 protein expression by western blot of whole cell lysates from GM-CSF-differentiated HMDM from healthy controls (HC) and DADA2 patients (P) untreated or treated with 5 μg/mL tunicamycin (TM) or 1 μg/mL brefeldin A (BFA) for 24 h. **(D)** $ADA2^{WT/WT}$ and $ADA2^{-/-}$ U-937 cells transduced with pathogenic ADA2 variants were incubated with or without 2.5 μg/mL tunicamycin for 24 h. ADA2 expression was measured by western blot compared with GM-CSF-differentiated HMDM from a healthy control (HC) and two DADA2 patients (P). **(E)** ADA2 protein expression in whole cell lysates and supernatant (SN) of U-937 and THP-1 cells left untreated or treated with 1 μg/mL tunicamycin (TM) or 1 μg/mL brefeldin A (BFA) for 24 h. (F) ADA2 secreted from M-CSF-differentiated human monocyte-derived macrophages from healthy control HC8 and DADA2 patient P8. ADA2 was detected using anti-ADA2 antibody clone EPR25430-131 (#ab288296; abcam) in all depicted blots. Triangles indicate HMW-ADA2 (dark blue) and LMW-ADA2 (light blue).

### Figure S3

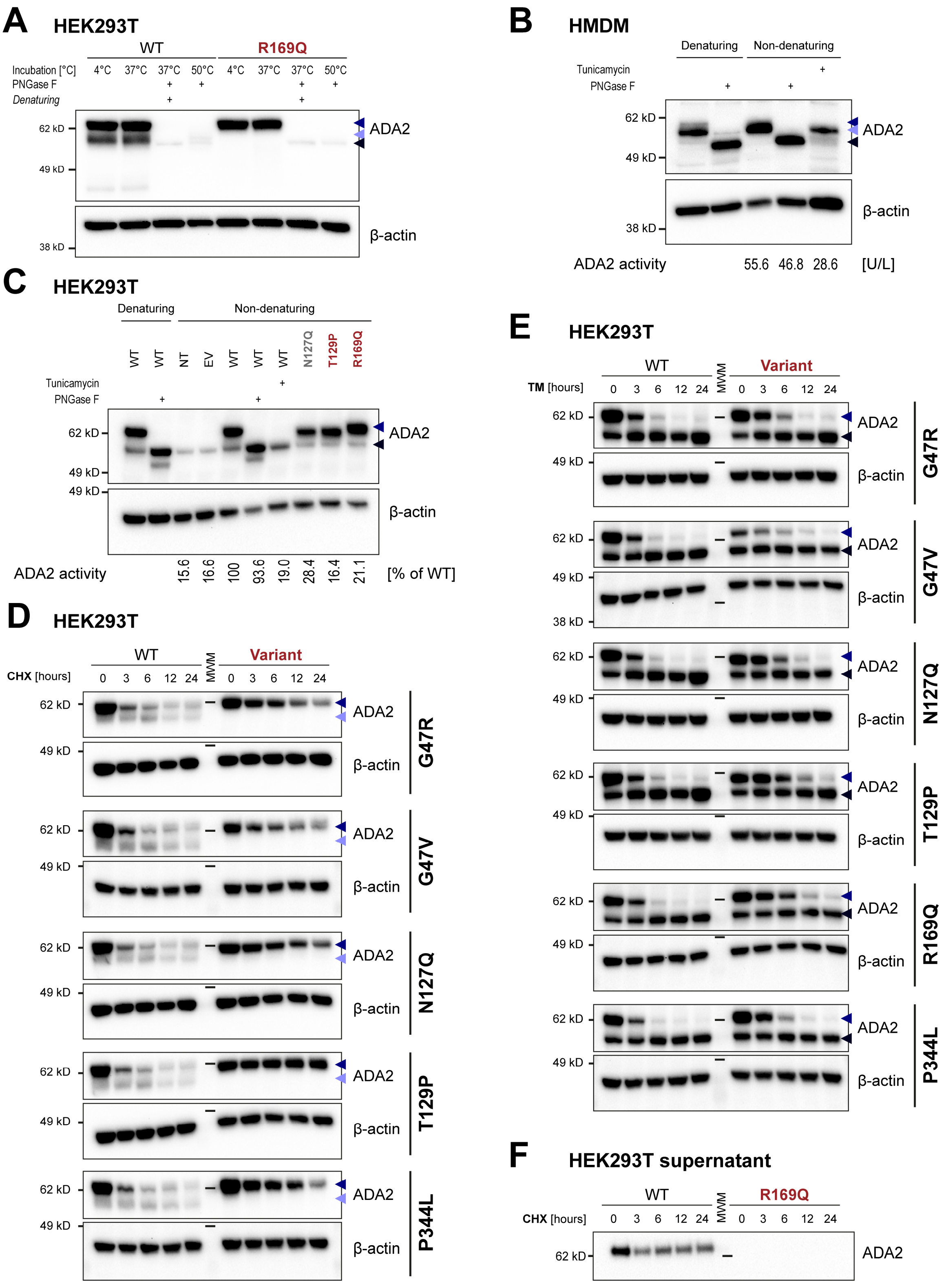

A
HEK293T
WT
R169Q
Incubation [°C]
4°C 37°C 37°C 50°C 4°C 37°C 37°C 50°C
PNGase F
Denaturing
62 kD
49 kD
38 kD
ADA2
β-actin
B
HMDM
Denaturing
Non-denaturing
Tunicamycin
PNGase F
62 kD
49 kD
38 kD
ADA2
β-actin
ADA2 activity
55.6 46.8 28.6
[U/L]
C
HEK293T
Denaturing
Non-denaturing
WT WT NT EV WT WT WT N127Q T129P R169Q
Tunicamycin
PNGase F
62 kD
49 kD
49 kD
ADA2
β-actin
ADA2 activity
15.6 16.6 100 93.6 19.0 28.4 16.4 21.1
[% of WT]
D
HEK293T
WT
Variant
CHX [hours]
0 3 6 12 24
MWM
0 3 6 12 24
62 kD
49 kD
ADA2
β-actin
G47R
G47V
N127Q
T129P
P344L
E
HEK293T
WT
Variant
TM [hours]
0 3 6 12 24
MWM
0 3 6 12 24
62 kD
49 kD
38 kD
ADA2
β-actin
G47R
G47V
N127Q
T129P
R169Q
P344L
F
HEK293T supernatant
WT
R169Q
CHX [hours]
0 3 6 12 24
MWM
0 3 6 12 24
62 kD
ADA2

**Figure S3: ADA2 expression upon modification of N-glycosylation (A)** Detection of glycan-free ADA2 by the anti-ADA2 antibody clone EPR25430-131. HEK293T cells were transfected with wild-type (WT) ADA2 or the pathogenic variant R169Q. Glycan removal of whole cell lysates was performed under denaturing conditions (#P0704; New England Biolabs) or under non-denaturing conditions (#A39245; Gibco) by incubation with PNGase F for 1 hour at 37 °C and 50 °C, respectively. GM-CSF-differentiated healthy control HMDM **(B)** or HEK293T cells transfected with wild-type (WT) *ADA2* or pathogenic *ADA2* variants **(C)** were treated with or without 5 µg/mL and 2.5 µg/mL tunicamycin for 24 h, respectively. Glycan removal was performed by incubation with PNGase F at 37 °C under denaturing (1 h) and non-denaturing (4 h) conditions. ADA2 enzyme activity was determined in whole cell lysates handled under non-denaturing conditions. **(D)** HEK293T cells were transfected with wild-type (WT) *ADA2* or different pathogenic variants and treated with 500 µg/mL cycloheximide (CHX) over 24 h. Whole cell lysates were generated at the indicated time points and immunoblotted for ADA2 expression. **(E)** HEK293T cells were transfected with wild-type (WT) *ADA2* or different pathogenic variants and treated with 2.5 µg/mL tunicamycin (TM) over 24 h. Whole cell lysates were generated at the indicated time points and immunoblotted for ADA2 expression. **(F)** HEK293T cells were transfected with wild-type (WT) ADA2 or the pathogenic variant p.R169Q and treated with 500 µg/mL cycloheximide (CHX) over 24 h. Supernatant was collected at the indicated time points and ADA2 levels were determined by western blot. ADA2 detection using anti-ADA2 antibody clone EPR25430-131 (#ab288296; abcam) in panels A, D and F and using anti-ADA2 (#HPA007888, Sigma-Aldrich) in panels B, C and E. Triangles indicate HMW-ADA2 (dark blue), LMW-ADA2 (light blue) and glycan-free ADA2 (black). Legend: MWM, molecular weight marker.

### Figure S4

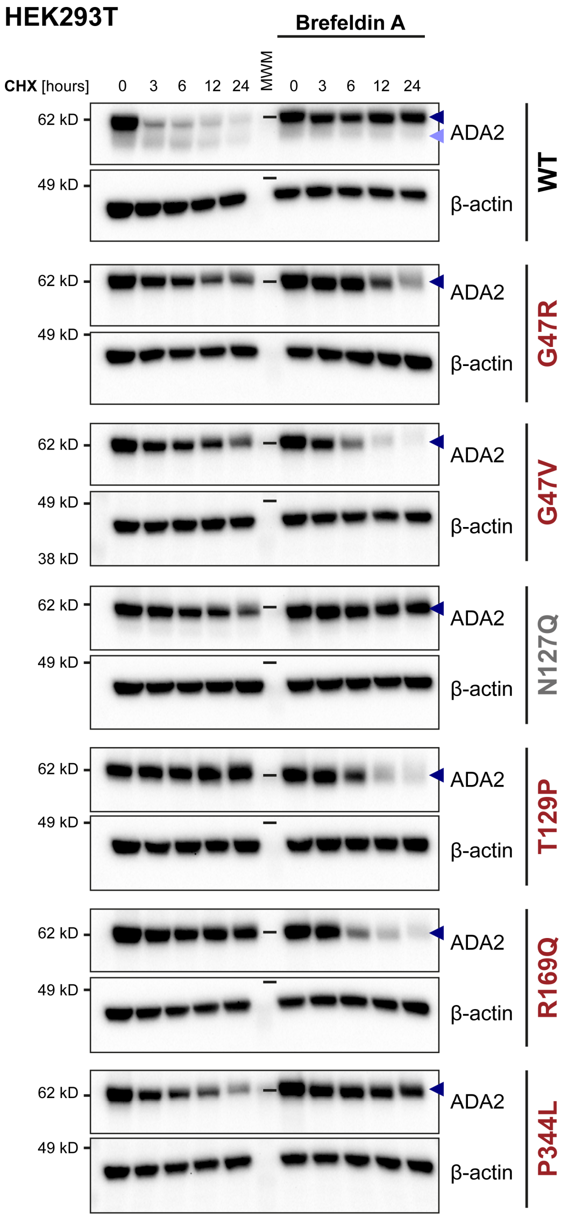


**Figure S4: ADA2 expression upon inhibition of ER to Golgi transfer.** HEK293T cells were transfected with wild-type (WT) *ADA2* or different pathogenic variants and treated with 500 µg/mL cycloheximide (CHX) ± 1 µg/mL brefeldin A over 24 h. Whole cell lysates were generated at the indicated time points and immunoblotted for ADA2 expression. ADA2 detection using anti-ADA2 antibody clone EPR25430-131 (#ab288296; abcam). Triangles indicate HMW-ADA2 (dark blue) and LMW-ADA2 (light blue). Legend: MWM, molecular weight marker.

### Figure S5

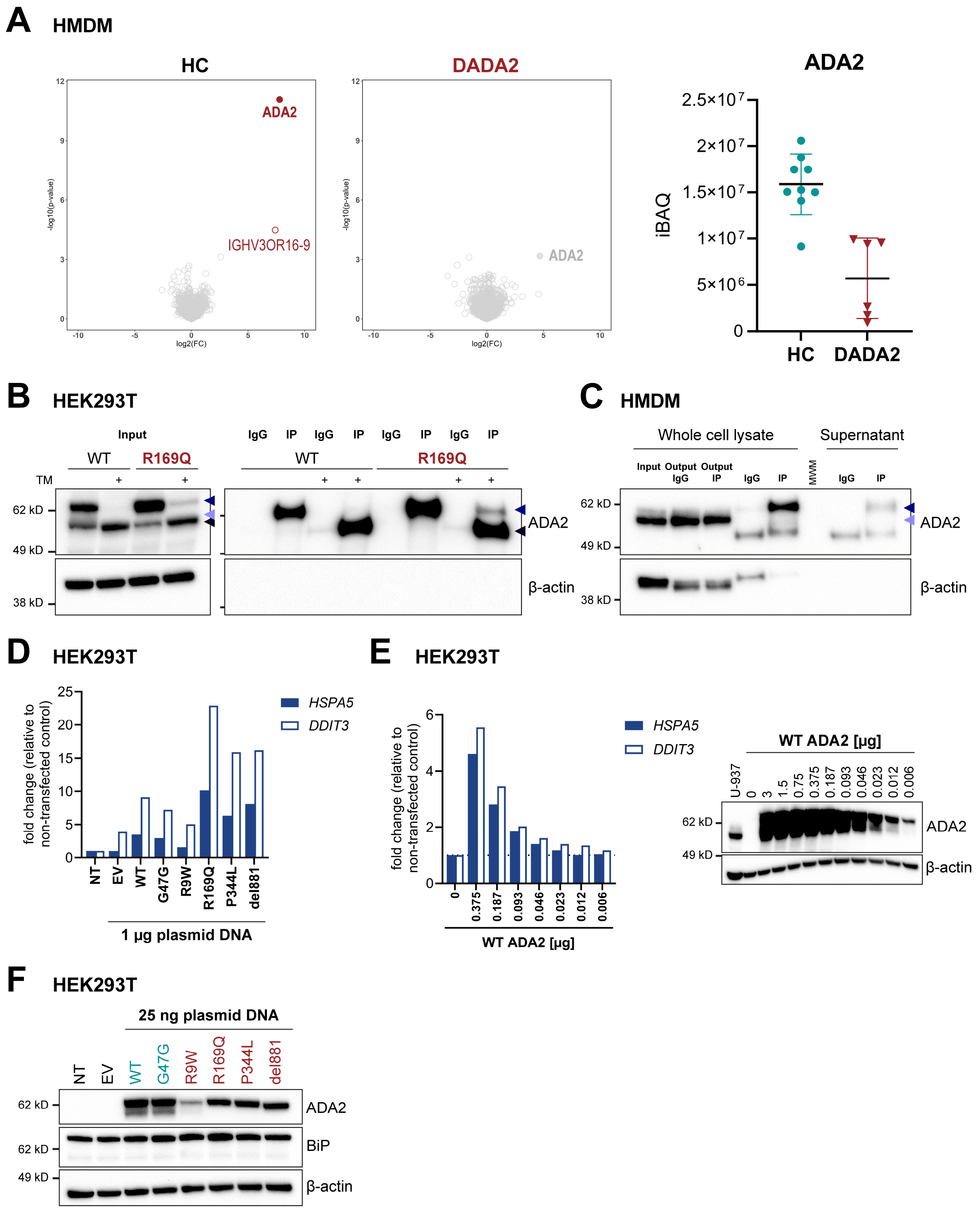

A HMDM
HC
ADA2
IGHV3OR16-9
-log10(p-value)
log2(FC)
DADA2
ADA2
ADA2
iBAQ
2.5×10^7
2×10^7
1.5×10^7
1×10^7
5×10^6
0
HC
DADA2
B HEK293T
Input
WT
R169Q
TM
IgG IP IgG IP IgG IP IgG IP
WT
R169Q
62 kD
49 kD
38 kD
ADA2
β-actin
C HMDM
Whole cell lysate
Supernatant
Input
Output IgG
Output IP
IgG
IP
MWM
IgG
IP
62 kD
49 kD
38 kD
ADA2
β-actin
D HEK293T
fold change (relative to non-transfected control)
HSPA5
DDIT3
NT EV WT G47G R9W R169Q P344L del881
1 µg plasmid DNA
E HEK293T
fold change (relative to non-transfected control)
HSPA5
DDIT3
0 0.375 0.187 0.093 0.046 0.023 0.012 0.006
WT ADA2 [µg]
WT ADA2 [µg]
U-937 0 3 1.5 0.75 0.375 0.187 0.093 0.046 0.023 0.012 0.006
62 kD
49 kD
ADA2
β-actin
F HEK293T
25 ng plasmid DNA
NT EV WT G47G R9W R169Q P344L del881
62 kD
62 kD
49 kD
ADA2
BiP
β-actin

**Figure S5: Immunoprecipitation of ADA2 and ER stress induction. (A)** $CD14^+$ monocytes from three healthy controls (HC) and two DADA2 patients (P) were differentiated into macrophages using 20 ng/mL GM-CSF for 10 days. Immunoprecipitation with the anti-ADA2 antibody clone EPR25430-131 (#ab288296; abcam) was followed by mass spectrometry. Binders significantly pulled down compared to isotype control with a false discovery rate of 10% are highlighted in red. The graph shows abundance of ADA2 in the IP samples of HC HMDM compared to DADA2 HMDM. The dot plots show median and interquartile range. **(B)** HEK293T cells transfected with wild-type (WT) ADA2 and the pathogenic variant p.R169Q were treated with or without 2.5 μg/mL tunicamycin (TM) for 24 h. ADA2 was pulled down from whole cell lysates using anti-DDK (FLAG) clone OTI4C5 (#TA50011, OriGene Technologies). Western blot detection was achieved using anti-ADA2 (#HPA007888, Sigma-Aldrich. **(C)** Western blot of ADA2 pulled down from the whole cell lysate (WCE) and supernatant (SN) of healthy control human monocyte-derived macrophages using anti-ADA2 antibody clone EPR25430-131 (#ab288296; abcam) for immunoprecipitation and detection with VeriBlot. All samples were prepared in four technical replicates. Three replicates were analyzed by mass spectrometry, the fourth replicate is shown on western blot. IgG samples represent isotype controls performed in parallel. The western blot is representative of n=5 (3 healthy controls; 2 patients). **(D)** HEK293T cells were transfected with 1 μg of plasmid DNA to overexpress wild-type (WT) *ADA2* and different pathogenic variants. The sham variant p.G47G was included as an internal control. Samples were taken 48h after transfection and ER stress was measured by mRNA expression of *HSPA5* and *DDIT3* normalized to *GAPDH*. Expression is depicted as $2^{-\Delta\Delta Ct}$ relative to non-transfected control (NT). Bar graphs show values of n=1 experiment. **(E)** Titration of plasmid DNA used for overexpression of ADA2. HEK293T cells were transfected with the indicated amounts of DNA to overexpress wild-type (WT) *ADA2*. Samples were taken 48h after transfection and ER stress was measured by mRNA expression of *HSPA5* and *DDIT3* normalized to *GAPDH*. Expression is depicted as $2^{-\Delta\Delta Ct}$ relative to the non-transfected control (0 μg). Bar graphs show values of n=1 experiment. **(F)** Western blot corresponding to the dot plots shown in Fig. 4B. ER stress response in HEK293T cells transfected with 25 ng wild-type (WT) *ADA2* and different pathogenic variants. Protein expression of BiP was determined 48h after transfection. The western blot is representative of n=6 independent experiments. ADA2 was detected using the anti-ADA2 antibody clone EPR25430-131 (#ab288296; abcam).

### Figure S6

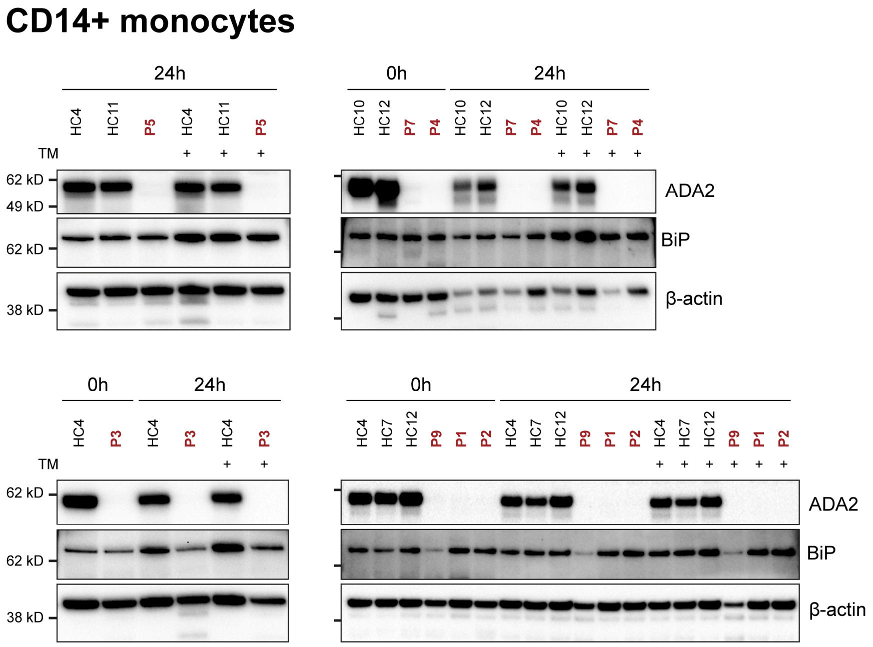


**Figure S6: BiP expression in CD14⁺ monocytes.** The depicted western blots correspond to the dot plots shown in Fig. 4C. ER stress response was evaluated by protein expression of BiP in whole cell lysates of $CD14^+$ monocytes from healthy controls (HC) and DADA2 patients (P) at baseline or after 24 h incubation with or without 1 µg/mL tunicamycin (TM). ADA2 was detected using the anti-ADA2 antibody clone EPR25430-131 (#ab288296; abcam).

### Figure S7

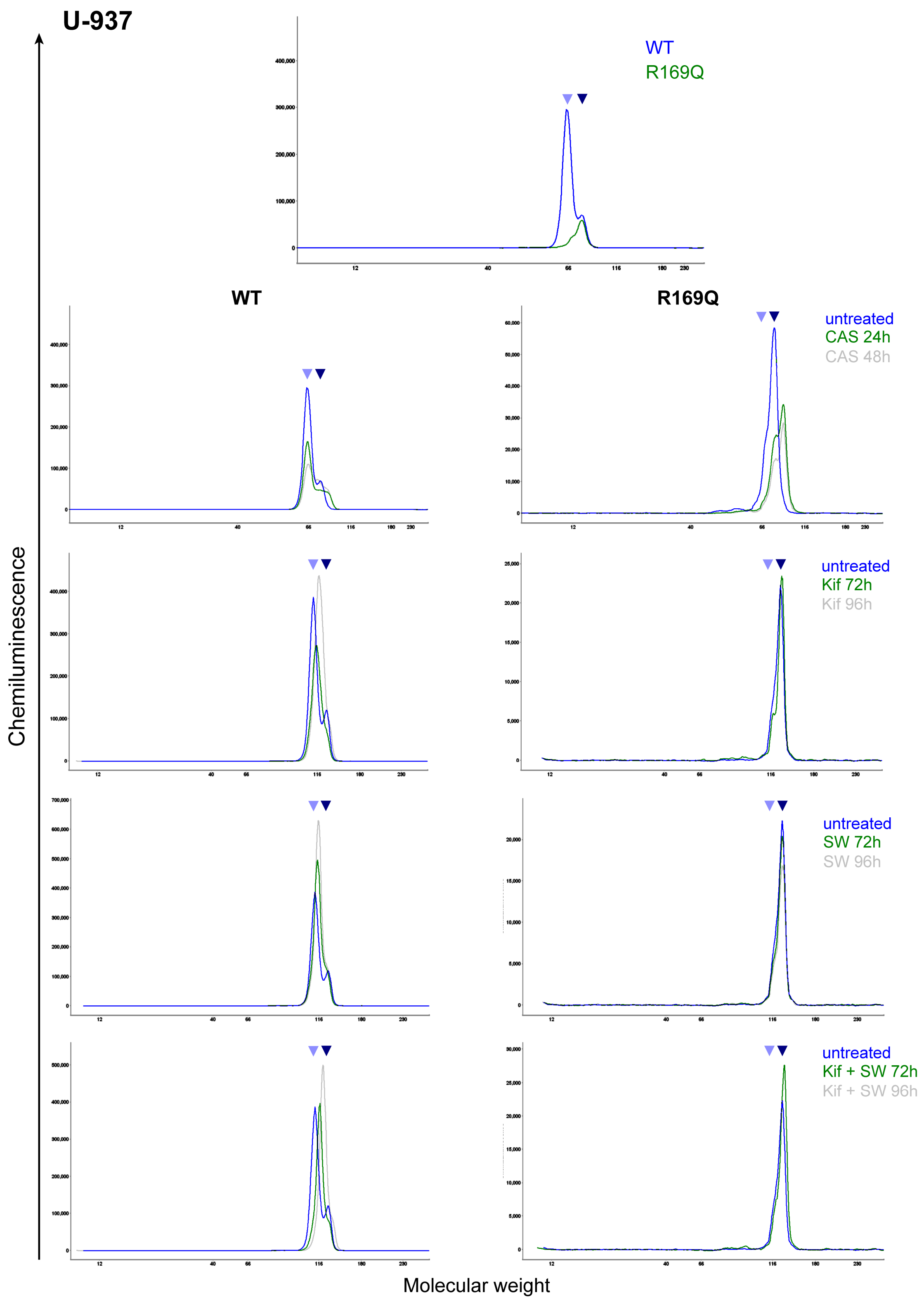
U-937
WT
R169Q
WT
R169Q
untreated
CAS 24h
CAS 48h
untreated
Kif 72h
Kif 96h
untreated
SW 72h
SW 96h
untreated
Kif + SW 72h
Kif + SW 96h
Chemiluminescence
Molecular weight

**Figure S7: Molecular weight of ADA2 after inhibition of glycan processing.** ADA2 protein was measured by Simple Western in ADA2$^{WT/WT}$ and ADA2$^{-/-}$ U-937 cells transduced with R169Q after 24 h or 48 h incubation with 100 µg/mL castanospermine (CAS) or after 72 h or 96 h incubation with 375 nM kifunensine (Kif) and/or 10 µM swainsonine (SW). The graphs show the signal for ADA2 by molecular weight. The top plot and third row left plot depict the original graphs shown in **Figure 6A**. ADA2 was detected using the anti-ADA2 antibody clone EPR25430-131 (#ab288296; abcam). Triangles indicate HMW-ADA2 (dark blue) and LMW-ADA2 (light blue).

### Figure S8

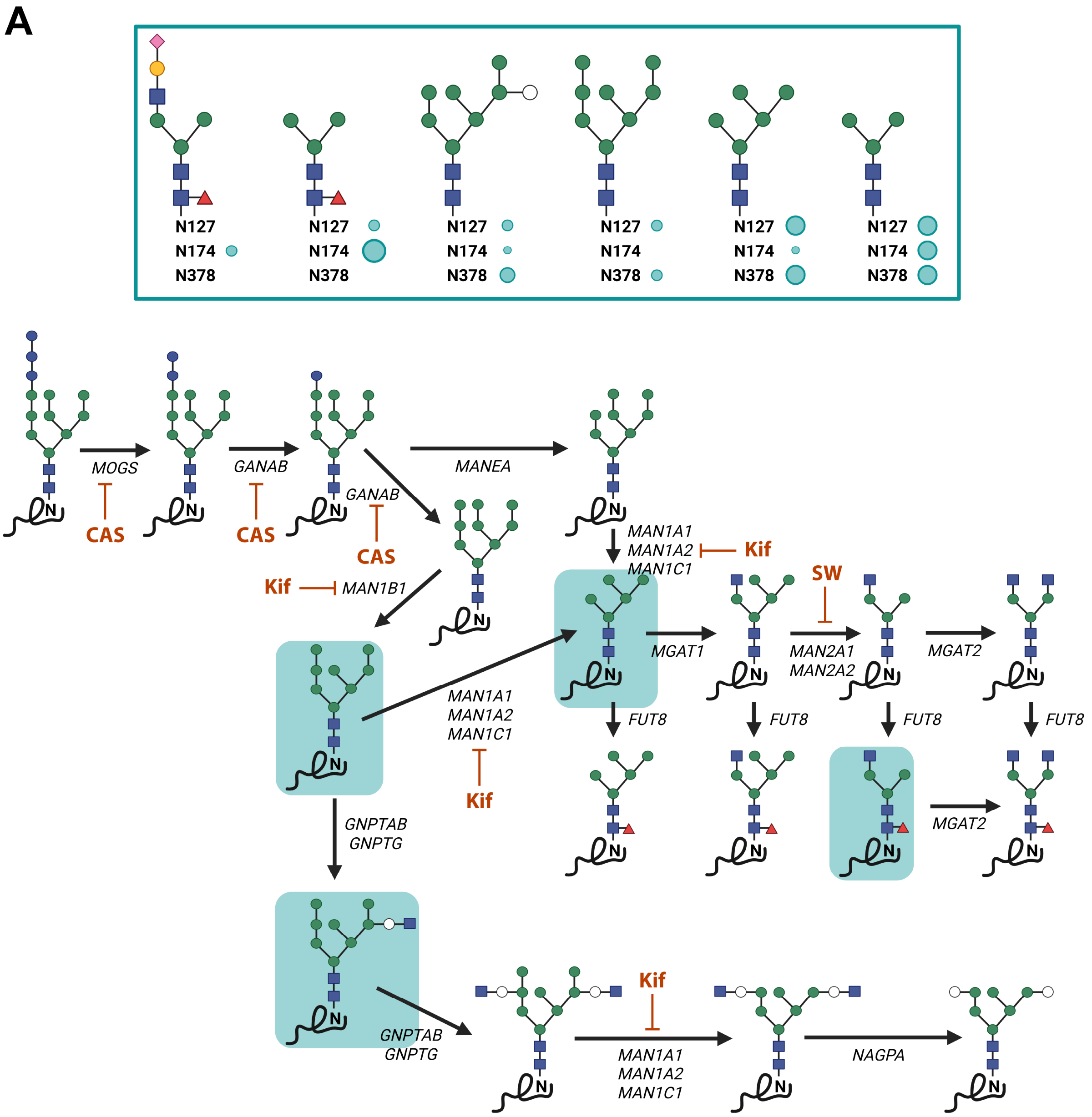
A
N127
N174
N378
MOGS
CAS
GANAB
CAS
GANAB
CAS
MANEA
MAN1A1
MAN1A2
MAN1C1
Kif
Kif
MAN1B1
SW
MGAT1
MAN2A1
MAN2A2
MGAT2
FUT8
FUT8
FUT8
FUT8
MAN1A1
MAN1A2
MAN1C1
Kif
GNPTAB
GNPTG
MGAT2
GNPTAB
GNPTG
Kif
MAN1A1
MAN1A2
MAN1C1
NAGPA

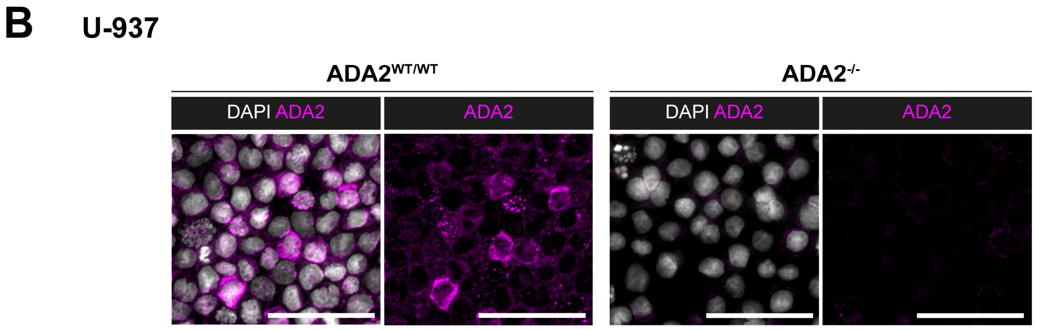
B
U-937
ADA2WT/WT
ADA2-/-
DAPI ADA2
ADA2
DAPI ADA2
ADA2

**Figure S8: Processing and imaging of ADA2. (A)** ADA2 glycan structures as shown by Greiner-Tollersrud et al. (*28*). The green circles indicate frequencies of the glycan trees at the respective glycosylation sites. Below, these glycan structures are shown in the context of the glycan processing pathways analyzed in this study. Carbohydrate residues are depicted according to the Symbol Nomenclature for Glycans (SNFG) (*20*). **(B)** Immunofluorescence microscopy of ADA2 in $ADA2^{WT/WT}$ and $ADA2^{-/-}$ U-937 cells to confirm staining specificity. Legend: CAS, castanospermine; Kif, kifunensine; SW, swainsonine.

### Table S3

| | **LMW-ADA2** | | **Intracellular enzyme activity** | | **Extracellular enzyme activity** | |
|---|---|---|---|---|---|---|
| ***Pearson correlation*** | ***r coefficient*** | ***P-value*** | ***r coefficient*** | ***P-value*** | ***r coefficient*** | ***P-value*** |
| **REVEL** | -0.611 | 2.6E-04 | -0.640 | 1.1E-04 | -0.709 | 1.7E-05 |
| **CADD** | -0.572 | 7.7E-04 | -0.570 | 8.2E-04 | -0.599 | 6.0E-04 |
| **SIFT** | 0.496 | 4.6E-03 | 0.540 | 1.7E-03 | 0.466 | 1.1E-02 |
| **PolyPhen** | -0.654 | 6.6E-05 | -0.639 | 1.1E-04 | -0.753 | 2.5E-06 |
| **Alpha missense** | -0.515 | 3.0E-03 | -0.590 | 4.7E-04 | -0.550 | 2.0E-03 |

**Table S3: Variant pathogenicity.** Correlation of LMW-ADA2 expression and enzymatic activity of *ADA2* variants with different pathogenicity scores. r represents Pearson's correlation coefficient.

### Table S1 (separate file)

**Table S1: Mass spectrometric analysis of top enriched proteins co-immunoprecipitated with ADA2 from HEK293T cells transfected with WT ADA2 or the variant p.R169Q.** Fold changes indicate pull-down by anti-DDK antibody targeting tagged ADA2 compared to isotype control.

### Table S2 (separate file)

**Table S2: Variant characteristics.** Pathogenicity scores were determined using the Ensembl Variant Effect Predictor. Allele frequencies were extracted from the gnomAD database. ADA2 enzyme activity is shown as previously reported by Jee et al. (*29*) and Sharma et al. (*44*).

#### Uncropped western blot images

The following pages display the uncropped images corresponding to the western blots presented in the main and supplementary figures of our manuscript.
